# Phylogenetic meta-analysis of chronic SARS-CoV-2 infections in immunocompromised patients shows no evidence of elevated evolutionary rates

**DOI:** 10.1101/2023.11.01.565087

**Authors:** Sanni Översti, Emily Gaul, Björn-Erik Ole Jensen, Denise Kühnert

## Abstract

Genomic sequences from rapidly evolving pathogens, sampled over time, hold information on disease origin, transmission, and evolution. Together with their sampling times, sequences can be used to estimate the rates of molecular evolution and date evolutionary events through molecular tip-dating. The validity of this approach, however, depends on whether detectable levels of genetic variation have accumulated over the given sampling interval, generating temporal signal. Moreover, different molecular dating methods have demonstrated varying degrees of systematic biases under different biologically realistic scenarios, such as the presence of phylo-temporal clustering.

Chronic SARS-CoV-2 infection in immunocompromised patients has been linked to remarkably higher intra-host molecular rates than those of global lineages, facilitating the emergence of novel viral lineages. Yet, most studies reporting accelerated rates lack the evaluation of temporal signal or comparison of multiple methods of inference, both required to reliably estimate molecular rates. In this study, we use 26 previously published longitudinally sampled sequence series obtained from chronically infected immunocompromised patients to re-evaluate the rate of SARS-CoV-2 intrahost evolution. Using a range of methods, we analyse the strength of temporal signal and infer evolutionary rates from tip-calibrated phylogenies. Regardless of heterogeneity in rate estimates between sample series and methods, we find within-host rates to be in good agreement with rates derived from host-to-host transmission chains.

Our findings suggest that when certain limitations of the methodology are disregarded, such as the underlying assumption of phylogenetic independence or the method’s sensitivity to phylo-temporal grouping, evolutionary rates can be substantially overestimated. We demonstrate that estimating within-host rates is a challenging question necessitating careful interpretation of findings. While our results do not support faster evolution across the complete viral genome during chronic SARS-CoV-2 infection, prolonged viral shedding together with relapsing viral load dynamics may nevertheless promote the emergence of new viral variants in immunocompromised patients.

**AUTHOR SUMMARY:** The evolutionary origin of SARS-CoV-2 variants of concern (VOC) is a longstanding point of controversy, with multiple proposed explanations. Observations of immunocompromised individuals being at a greater risk of developing a prolonged SARS-CoV-2 infection have led to the ‘Chronic infection hypothesis’, suggesting that these cases may contribute to the emergence of VOCs. Correspondingly, many studies have reported accelerated viral evolution of SARS-CoV-2 within immunocompromised individuals with respect to the viral background population. However, many of these findings have not been validated with appropriate analytical methods. In this study we re-evaluate the rate of intrahost viral evolution of SARS-CoV-2 within immunocompromised patients utilising a range of methods. We assess the performance of different methodologies and compare our results to published estimates of SARS-CoV-2 evolutionary rates. Our systematic comparison showed no evidence supporting the previous claims of elevated levels of intrahost evolution in immunocompromised patients with chronic SARS-CoV-2. Instead, our findings exemplify the complexity of within-host viral dynamics, suggesting that a more comprehensive understanding of SARS-CoV-2 evolutionary processes would be derived from concurrent evaluation of viral genomic data together with patients’ clinical information.

## INTRODUCTION

Molecular dating postulates that differences between two sequences are directly proportional to the time elapsed since they diverged [1], hence allowing an estimation of the timing of evolutionary events. Calibration of a molecular clock with independent temporal information is required to convert relative divergence times of a phylogenetic tree into absolute timescales. For serially sampled data sets, including those generated for rapidly evolving pathogens such as severe acute respiratory syndrome coronavirus 2 (SARS-CoV-2), trees can be calibrated using the sampling times of genetic sequences [2,3] (for review see [4]).

Whilst time stamped genealogies have become fundamental for understanding pathogen evolution, the accuracy of estimated evolutionary rates substantially influences the reliability of inferred time-scales (for definitions and discussion of different rates of evolution, see [5,6]). As a result, a large range of evolutionary models and methods have been developed, key distinctions between different methodologies relying on whether the method accommodates phylogenetic uncertainty and if rate heterogeneity amongst lineages can be modelled. In the simplest approach, a linear regression is fitted between sampling dates and corresponding root-to-tip genetic distances [7,8]. In spite of root-to-tip (RTT) regression analysis being extensively used, its assumptions of statistical independence of the sequences and rate homogeneity among lineages can be considered as substantial limitations [4,9,10]. Alternatively, least-squares dating (LSD) is another widely used distance-based approach which provides estimations of evolutionary rates determined by maximising the likelihood of the rooted phylogeny [11]. Whereas LSD has been demonstrated to be somewhat robust to rate heterogeneity [11], the evolutionary patterns of most datasets are more accurately described by relaxing the assumption of strictly clock-like evolution (for review see for example [12]). In response, distance-based phylogenetic approaches, such as TreeDater [13], have been implemented to explicitly account for branch specific evolutionary rates. Whereas all aforementioned distance-based methods rely on user-supplied fixed tree topology facilitating only the estimation of the root placement, probabilistic models implemented in a Bayesian framework can be used for joint estimation of phylogenetic tree topology and evolutionary rates (for an introduction on Bayesian phylogenetic analysis, see for example [14,15]). Due to their broad applicability, Bayesian phylogenetic methods, such as BEAST2 [16] and RevBayes [17], have become widely utilised for molecular dating. In addition to tree uncertainty these methods can accommodate complex demographic and evolutionary models, such as an uncorrelated relaxed clock model where rate associated with each branch is independently drawn from a shared underlying distribution [18].

Irrespective of the phylogenetic approach chosen, a prerequisite for molecular dating analysis of tip-calibrated phylogenies is that genetic changes can be considered to have accumulated rapidly enough relative to the available range of sequence sampling times. If measurable levels of genetic variation have accumulated over a given sampling interval, the population is considered as *‘measurably evolving’* [8]. Since insufficient temporal signal might lead to biased estimates of rates and timescales, determining the strength of temporal signal of heterochronously sampled data is an essential step prior to the estimation of evolutionary rates [19]. As a simplest interpretation of adequate temporal signal can be considered a positive correlation between sequence sampling times and their corresponding root-to-tip distances (see for example Fig.2 in [4]). However, since RTT can be viewed as a qualitative method that only provides visual evidence for sufficient temporal signal [9], more sophisticated approaches, such as the ‘Date-randomization test’ (DRT, [20]) and ‘Bayesian Evaluation of Temporal signal’ (BETS, [21]), have been developed for enhanced temporal signal assessment.

Since the onset of the coronavirus disease 2019 (COVID-19) pandemic, tip-calibrated phylogenies have been exploited extensively to gain insights into the origin and spread of SARS-CoV-2 (for review see [22]). Despite within-patient viral genetic diversity appearing to be quite limited over the duration of an acute infection [23–25] rather soon after the initial outbreak, the virus exhibited a significant number of genetic differences through time [10]. Consequently, a wealth of studies has estimated evolutionary rates for SARS-CoV-2 at the population level yielding mean estimates between 5.75e-04 and 1.60e-03 substitutions/site/year converting to ∼1.4–4.0 substitutions per genome per month (see Table 1 in [22]). Whereas molecular dating approaches have been used rather routinely to infer molecular rates of between-host transmission chains, their full potential has not yet been exploited to evaluate intrahost evolution of SARS-CoV-2. In contrast to the majority of infected individuals with viral load cleared generally from 10 to 16 days after the onset of symptoms [26,27], numerous independent studies have shown that immunocompromised individuals with diverse clinical backgrounds are at greater risk of developing a prolonged SARS-CoV-2 infection (for references, see Table 1). This long-term viral shedding might provide favourable conditions for intrahost viral evolution [28,29] facilitating emergence of new variants that consequently can transmit to the general population. Accordingly, the *‘Chronic Infection Hypothesis’* [30] states that prolonged infections in immunocompromised patients have shaped the evolution of SARS-CoV-2 by acting as a source of variants of concern. In agreement with the proposed hypothesis a large number of studies have reported accelerated SARS-CoV-2 evolution within immunocompromised individuals, suggesting up to two-fold higher molecular rates when accounting for the whole SARS-CoV-2 genome [30–37].

**Table 1.** Overview of sample series included in this study. * Defined with Nextclade v2.14.1. ** For details, see supplementary table S2.

| Sample series | Number of sequences included in the analysis | Sampling window (days) | Pango lineage * | Patient's underlying clinical condition ** | Reference |
| --- | --- | --- | --- | --- | --- |
| Baang-pt-1 | 9 | 99 | B.1.576 | B-cell neoplasm | [44] |
| Brandolini-pt-1 | 8 | 86 | AY.122 | B-cell neoplasm | [37] |
| Caccuri-pt-1 | 12 | 222 | B.1.1 | B-cell neoplasm | [45] |
| Chaguza-pt-1 | 30 | 392 | B.1.517 | B-cell neoplasm | [30] |
| Choi-pt-1 | 9 | 134 | B.1.576 | Rheumatological/<br>autoimmune disease | [31] |
| Ciuffreda-pt-1 | 16 | 129 | A.2 | PID | [32] |
| Gandhi-pt-1 | 15 | 141 | B.1.576 | B-cell neoplasm | [46] |
| Halfmann-pt-1 | 12 | 373 | B.1.2 | B-cell neoplasm<br>and PID | [47] |
| Harari-pt-5 | 9 | 75 | B.1.1.50 | B-cell neoplasm | [41] |
| Huygens-pt-2 | 13 | 160 | BA.1.1 | B-cell neoplasm | [48] |
| Jensen-pt-2 | 8 | 22 | B.1.1 | HIV/AIDS | [49] |
| Kemp-pt-1 | 16 | 100 | B.1.1.1 | B-cell neoplasm | [39] |
| Khatamzas-pt-1 | 21 | 149 | B.1.1 | B-cell neoplasm | [50] |
| Lee-pt-11 | 11 | 64 | B.1 | B-cell neoplasm | [51] |
| Lee-pt-4 | 8 | 342 | B.1.576 | B-cell neoplasm | [51] |
| Li-pt-1 | 10 | 140 | B | Multiple clinical<br>conditions | [43] |
| Lynch-pt-1 | 8 | 77 | B.1.1 | B-cell neoplasm | [52] |
| Pérez-Lago-pt-1 | 9 | 123 | B | B-cell neoplasm | [40] |
| Pérez-Lago-pt-2 | 10 | 117 | B.1 | B-cell neoplasm | [40] |
| Riddell-pt-2 | 9 | 111 | B.1.1.7 | B-cell neoplasm and<br>HIV/AIDS | [53] |
| Riddell-pt-3 | 15 | 255 | B.1.1.7 | HIV/AIDS | [53] |
| Rockett-pt-2 | 8 | 31 | AY.39.1.2 | PID | [54] |
| Rockett-pt-4 | 8 | 40 | AY.39.1.3 | Myelodysplastic<br>syndrome/<br>myeloproliferative<br>disorder | [54] |
| Rockett-pt-8 | 12 | 34 | AY.39.1 | B-cell neoplasm and<br>multiple other clinical<br>conditions | [54] |
| Sonnleitner-pt-1 | 10 | 98 | B.1.1.232 | B-cell neoplasm | [38] |
| Weigang-pt-1 | 9 | 140 | B.1.1 | Multiple clinical<br>conditions | [55] |

Intriguingly, while it has been asserted in a number of publications that intrahost evolutionary rates in immunocompromised patients are noticeably higher, most often the findings are not being supported by any substantive analytical method. Instead, most commonly reported rates are determined by directly calculating the number of mutations accumulated [32,34,38] or through root-to-tip regression analysis [30,36]. While the latter’s limitations have already been discussed, the former may result in an overestimation of the number of changes due to general assumption of changes accumulating over time in a single viral lineage. This contradicts observations of within-host SARS-CoV-2 viral populations being frequently a collection of genetically closely related lineages, i.e. coexisting quasispecies [30,37,39–41]. Furthermore, no comparison of different molecular dating methods has been performed, nor the degree to which they might be relied upon has been tested. More importantly, the strength of the temporal signal of within-host sample series has not been evaluated, leaving the conclusions rather speculative. However, as prolonged SARS-CoV-2 infections within immunocompromised individuals have supposedly played a key role in shaping the COVID-19 pandemic, compelling interests for public health exist to understand more thoroughly the interplay between chronic SARS-CoV-2 infection and viral evolution.

In this study, we re-evaluate the rate of intrahost molecular evolution of SARS-CoV-2 within chronically infected immunocompromised patients. Our dataset consists of previously published SARS-CoV-2 sequences sampled from 26 patients at multiple time points over the course of infection. For each sample series, we evaluate the strength of the temporal signal and subsequently infer evolutionary rates based on tip-calibrated phylogenies using a variety of methods – including distance-based methods as well as Bayesian inference. We evaluate the performance of methods used and compare the results with earlier estimates. Through this systematic assessment our aim is to bring into the awareness of researchers aiming to infer within-host evolutionary dynamics through molecular dating the important limitations of some of the approaches used. Our results show that by ignoring the evaluation of temporal signal and the constraints of the phylogenetic method used, inferred evolutionary rate estimates may be substantially distorted, while actual patterns of viral evolution may go undiscovered. Therefore, we propose that the framework developed in this study should be considered in future studies utilising phylogenetic inference to infer intrahost molecular rates. Furthermore, we explore novel methods of combining phylogenetic inference with published clinical metadata. Whereas our results in general do not lend support for accelerated intrahost viral evolution of SARS-CoV-2 across the complete viral genome, prolonged viral shedding together with the relapsing viral load dynamics may nevertheless promote the emergence of novel viral variants, such as variants of concern.

## RESULTS

Heterochronously sampled sequence series from immunocompromised patients were used to re-evaluate SARS-CoV-2 intrahost evolutionary rates over the course of chronic viral infection. Sample series were identified through a literature search and for all datasets genetic diversities were determined. Preliminary assessment of temporal signal was performed with RTT followed by evolutionary rate estimation with LSD2 and TreeDater. For those sample series, for which evidence of temporal signal was considered sufficient, rates were further inferred with Bayesian inference. For a subset of sample series, we additionally evaluated the temporal simultaneity of the changes in evolutionary rate across phylogenetic branches with changes in viral load dynamics and the timing of SARS-CoV-2 treatments administered (‘Patient case histories’). A schematic overview of the workflow is presented in Figure 1.

**Figure 1.**
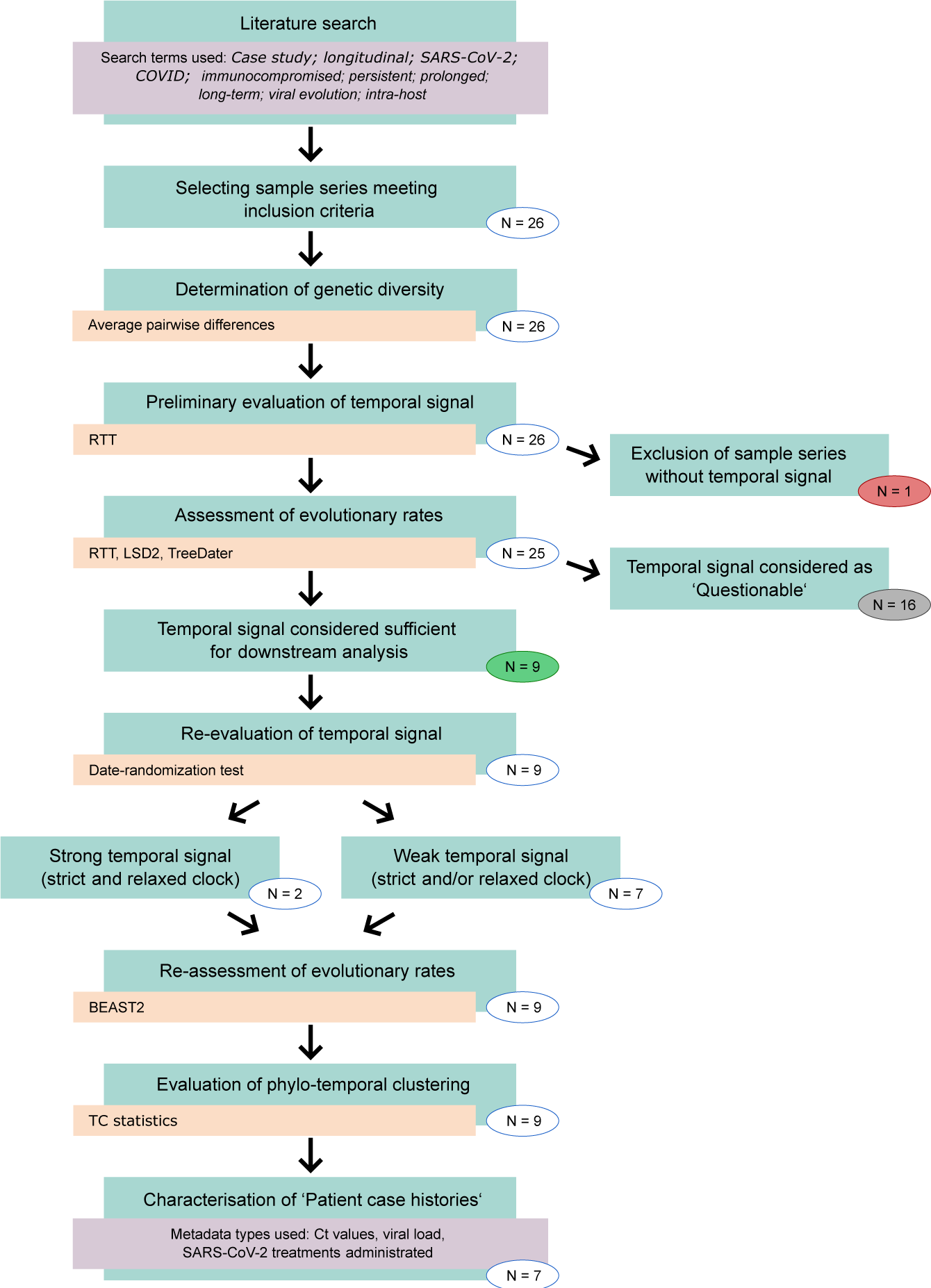
Schematic overview of the workflow. Number of sample series included in each step are given within the circles. Colouring of the number of the sample series corresponds to Figure 2 (red = no temporal signal, grey = questionable temporal signal, green = sufficient temporal signal). Software/Method or statistics used are indicated with yellow boxes. Additional information is indicated with purple boxes.

**Figure 2.**
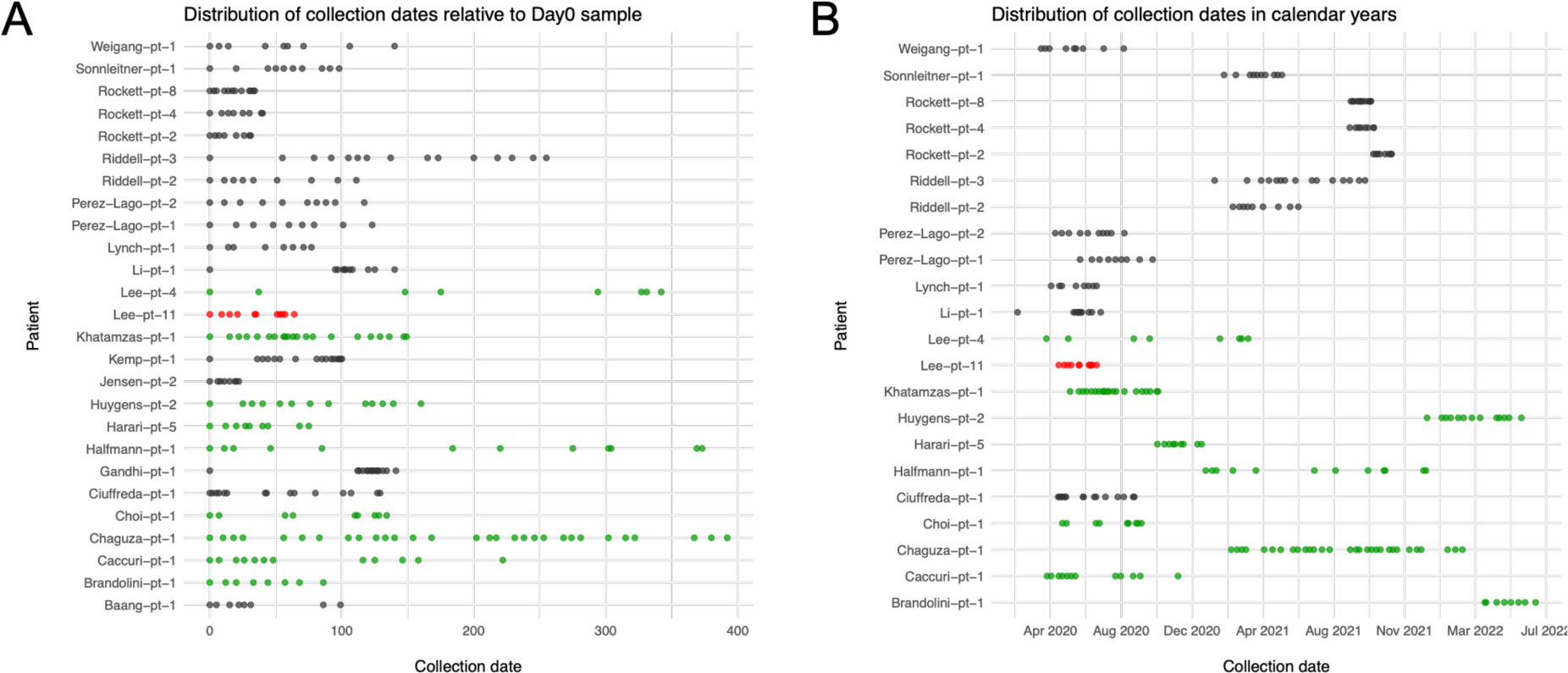
Temporal distribution of sample collection points. In figure 2A collection dates are given relative to the first sample of each sample series (Day0) whereas in figure 2B collection dates are represented in calendar years. Sample series are colour coded according to their temporal signal: Red indicates patients with no temporal signal, grey indicates poor (‘Questionable’) temporal signal whereas green denotes patients with sufficient temporal signal (evaluated based on analysis with RTT, LSD2 and TreeDater). As for the following patients the collection dates were not given as calendar units, they are omitted from figure 2B: Baang- pt-1, Gandhi-pt-1, Jensen-pt-2 and Kemp-pt-1.

### Data Collection

All data analysed during this study were obtained through a literature search, resulting in the identification of 85 publications presenting chronic SARS-CoV-2 sample series (for details, see Materials & Methods). In order to minimise the phylogenetic uncertainty and thus increase the precision of evolutionary rate estimates, we chose to include only sample series for which eight or more viral consensus sequences from unique collection dates were available. Additionally, inclusion criteria required evidence in the original publication confirming the immunocompromised status of the patient and the occurrence of a long-term infection, hence excluding multiple independent infections or superinfection. Following the procedure presented in [41] an individual was considered to have a chronic SARS-CoV-2 infection if there was evidence of sustained high viral loads for a period of at least 20 days. In total, 26 patients met all criteria. Clinical metadata and sequence accession information are reported within the Supplementary Materials (Supplementary tables S1–S7).

The final data, comprising 304 sequences from 26 patients, included one sample obtained from the gastrointestinal tract (Kemp-pt-1 Day85 stool sample) and one sample obtained from serum (Pérez-Lago-pt-2 Day40). The remaining samples were derived from the respiratory tract including nasopharyngeal, oropharyngeal, combined nasopharyngeal/oropharyngeal, sputum, bronchoalveolar lavage and tracheal aspirate specimen types. The number of sequences per sample series varied from eight up to 30 sequences (Table 1). The sampling windows, i.e. the days between the first and last sequence sampling point for each sample series, ranged from 22 days (Jensen-pt-2) to 392 days (Chaguza-pt-1) (Table 1, Figure 2A). Collection date information was available in calendar units for 22 sample series and altogether these covered a time period from February 2020 to June 2022 (Figure 2B). Sample series represented in total 16 different Pango lineages [42]. Lineages B, B.1, B.1.1, B.1.1.7 and B.1.576 were observed more than once (Table 1, Supplementary table S8). Seven of the patients carried lineages identified as variants of concern: Alpha (Riddell-pt-2 and Riddell-pt-3), Delta (Brandolini-pt- 1, Rockett-pt-2, Rockett-pt-4 and Rockett-pt-8) and Omicron (Huygens-pt-2) (Supplementary table S8). The assignment of Pango lineages to Li-pt-1 sample series with Nextclade version v2.14.1 suggested that samples reflected distinct lineages (Supplementary table S1). However, since the original paper [43] regarded strong sequence similarity as evidence against reinfection, we decided to include the sample series in the subsequent study. Nevertheless, results should be interpreted with caution.

18 of the patients were receiving treatment for B-cell neoplasm (including B-cell lymphoma and B-cell leukemia), 3 each for primary immunodeficiency (PID) and for HIV/AIDS, 1 for myelodysplastic syndrome/myeloproliferative disorder and 1 for rheumatological/autoimmune disease as well as 3 patients with other forms of immunodeficiency (Table 1). Some of the patients had more than one disease associated with immunodeficiency (Supplementary table S2). Due to highly unequal representation of distinct underlying clinical condition categories, analytical comparisons between categories were not feasible. Therefore, the potential differences in how different underlying clinical conditions may influence the intrahost evolution of SARS-CoV-2 were not further explored nor discussed within this study.

### Assessment of genetic diversity among sample series and temporal signal with RTT

To approximate the genetic diversity of each sample series we determined the mean number of pairwise differences between sequence pairs within each dataset (Supplementary table S9). Whereas approximately half of the sample series displayed low levels of genetic diversity with observed mean pairwise differences being less than 5.0, for some of the sample series differences were notably higher, yielding mean values above 10.0. As we detected genetic changes within all sample series, the strength of the temporal signal was firstly assessed with the regression of root-to-tip distances (RTT). RTT indicated a positive correlation between the genetic root-to-tip distances and the sampling times for all sample series (Supplementary figure S1). However, assuming that the strength of the temporal signal can be evaluated based on RTT plots and correlation coefficient values, the sample series displayed highly variable levels of temporal signal, with R^2^ values ranging between 0.23 and 0.99. The low R^2^ value of 0.23 and p-value of 6.45e-02 observed for sample series Lee-pt-11 was considered to indicate inadequate temporal signal, and we chose to exclude this data from subsequent analyses. For all the remaining sample series p-values were below the assumed threshold of 0.05, despite the R^2^ values being rather low, Riddell-pt-3 displaying the lowest value of 0.39 (Supplementary figure S1). Based on positive correlation between genetic differences and sequence sampling dates observed alone, 25 of the sample series included in this study would be suitable for phylogenetic molecular clock analysis [9]. However, subsequent analyses with LSD2 and TreeDater excluded many of these, showing adequate temporal signal for only nine sample series (Figure 2). For the remaining 16 sample series a lack of sufficient temporal signal was detected and therefore the temporal signal was considered as ‘Questionable’ (for details, see Methods). Among these 16 datasets, for one dataset the rate estimate was successfully determined only with RTT (Rockett-pt-4) whereas for three sample series estimates were obtained with RTT and TreeDater but not with LSD2 (Riddell-pt-3, Sonnleitner-pt-1 and Weigang-pt-1).

The majority of the sample series for which LSD2 and TreeDater exhibited poor performance displayed rather low genetic diversities (i.e. Baang-pt-1, Jensen-pt-2, Lynch-pt-1, Pérez-Lago- pt-1, Pérez-Lago-pt-2, Riddell-pt-2, Riddell-pt-3, Rocket-pt-2, Rocket-pt-4, Weigang-pt-1) (Supplementary table S9). For some sample series with higher diversity the absence of strong temporal signal might be explained by highly skewed temporal distributions of sampling points (i.e. Gandhi-pt-1, Kemp-pt-1 and Li-pt-1, Figure 2). Genetic diversities for sample series with questionable and sufficient temporal signals showed positive correlations between sampling windows with correlation coefficient values of R^2^=0.42 and R^2^=0.84 (Figure 3). However, for sample series with questionable temporal signal correlation was not statistically significant (p=0.11). This indicates that the duration of infection can explain only some of the observed genetic diversity, meaning that novel mutations emerge with highly variable patterns among sample series.

**Figure 3.**
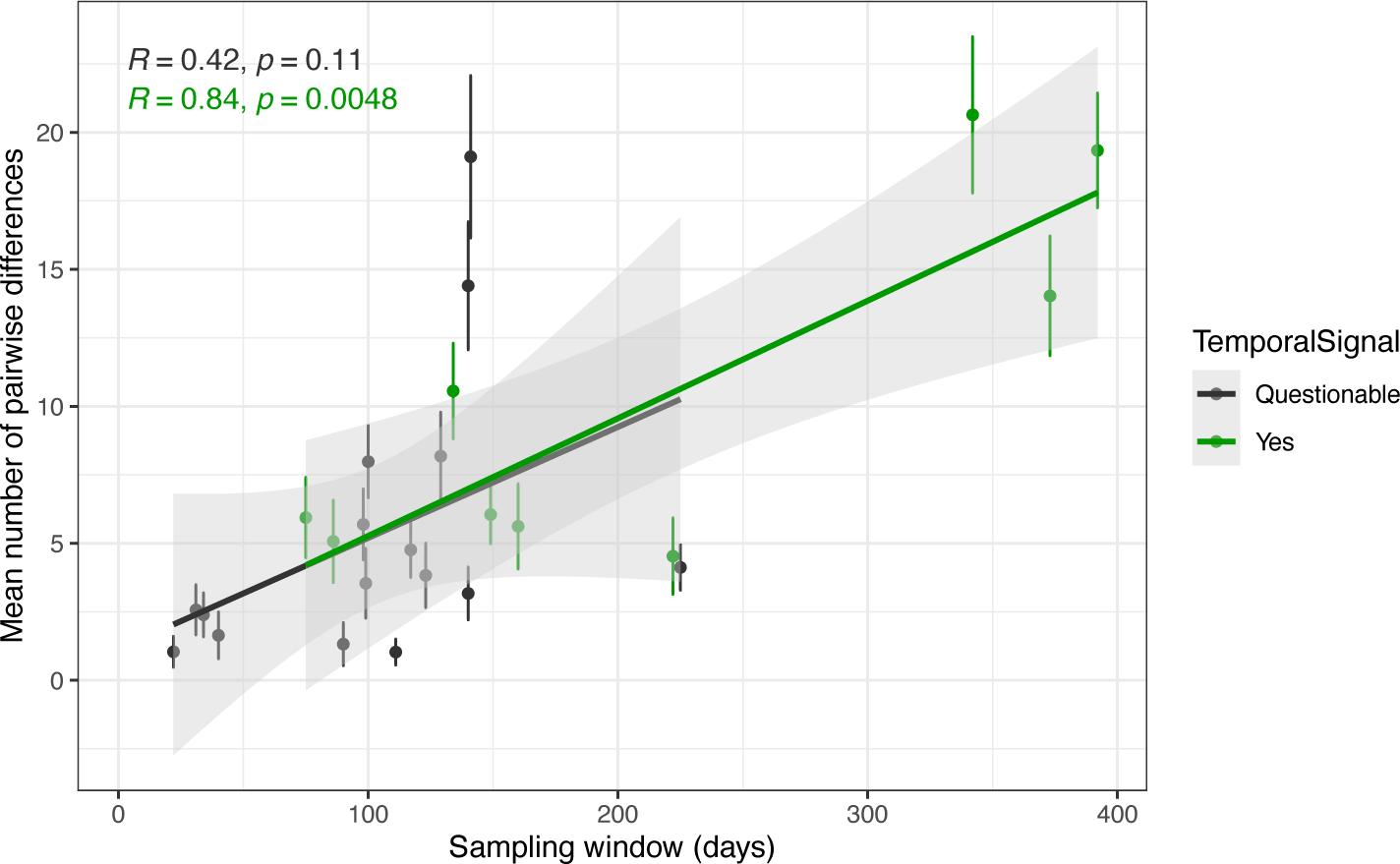
Mean number of pairwise differences between sequence pairs within each sample series plotted against the sampling window. Circles represent mean estimates and vertical lines standard deviations for each sample series. Green colour denotes sample series for which temporal signal was considered sufficient based on LSD2 and TreeDater analysis whereas grey colour denotes sample series for which temporal signal was not adequately assigned. Based on a linear regression model statistically significant indications of strong correlations between sampling window and mean number of pairwise distances was found only for a group of sample series exhibiting adequate temporal signal.

### Evolutionary rate estimates – RTT, LSD2 and TreeDater

Comparison of evolutionary rates obtained with RTT, LSD2 and TreeDater, reveals notable discrepancies across the estimates between different sample series as well as between different methods (Figures 4 and 5). Inconsistencies among methods were observed for sample series with and without adequate temporal signal. For the nine sample series with sufficient temporal signal LSD2 and TreeDater yielded comparable mean rate estimates within each dataset (Figure 4, Supplementary table S10). Estimates obtained with RTT were consistently higher than either of these. A similar pattern of elevated RTT estimates was seen also for the 15 sample series with modest temporal signal (Figure 5, Rockett-pt-4 excluded as no estimates were obtained with LSD2 nor with TreeDater). Within each dataset no significant differences were detected between estimates produced with TreeDater by assuming strict or relaxed clock models. Similarly, LSD2 produced highly congruent estimates with and without collapsing the short branches of the tree. For LSD2 we additionally evaluated the possible impact of an outgroup inclusion and re-inferred rate estimates for trees rooted with the SARS-CoV-2 reference sequence (GenBankID: NC_045512.2 [56]). As shown in Supplementary figure S2, usage of an outgroup taxon did not have a significant impact on the inferred rates.

**Figure 4.**
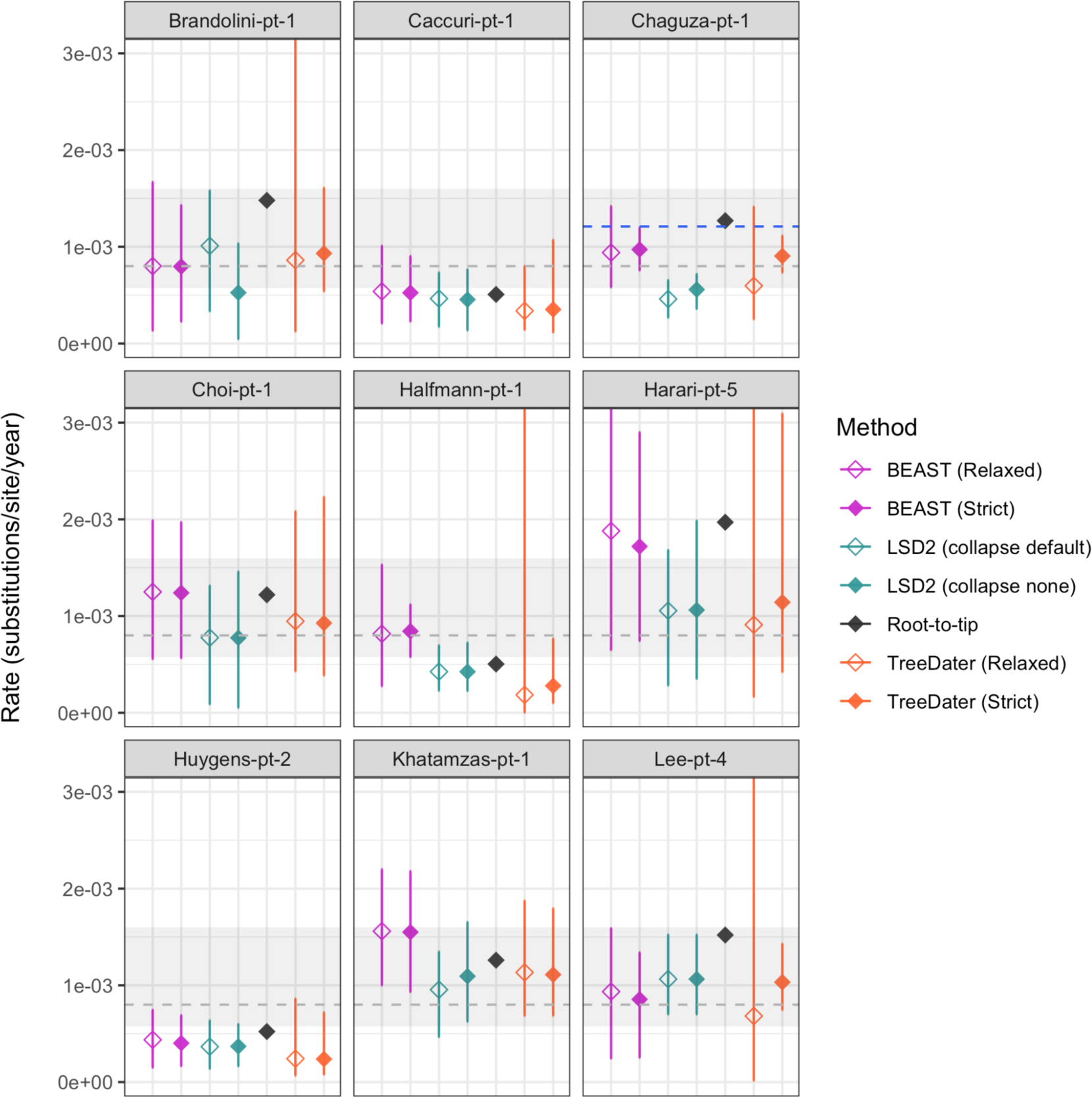
Evolutionary rate estimates determined for nine patients with sufficient temporal signal (strength of temporal signal defined here based on RTT as well as TreeDater and LSD2 results). For all sample series, rates were determined with following methods: BEAST2 (by assuming relaxed and strict clock models), LSD2 (with and without collapsing short branches), root-to-tip and TreeDater (by assuming relaxed and strict clock models). In each panel, the Y axis denotes the evolutionary rate in substitutions/site/year. For estimates inferred with BEAST, diamonds represent median estimates and associated vertical lines correspond to 95% highest posterior density intervals (HPDI). For RTT only point estimates are represented. For other distance-based methods (i.e. LSD2 and TreeDater) diamonds represent mean estimates and bars illustrate confidence intervals. Grey dashed line represents the commonly used SARS-CoV-2 substitution rate estimate of 8.00e-04 substitutions/site/year. The grey shaded area denotes the lowest and highest mean evolutionary rate estimates for SARS-CoV-2 collected from various publications (5.75e-04 – 1.60e-03 subst./site/year, see Supplementary table S11). For Chaguza-pt-1 a previous estimate of 1.2e-03 substitutions/site/year is indicated with a blue dashed line.

**Figure 5.**
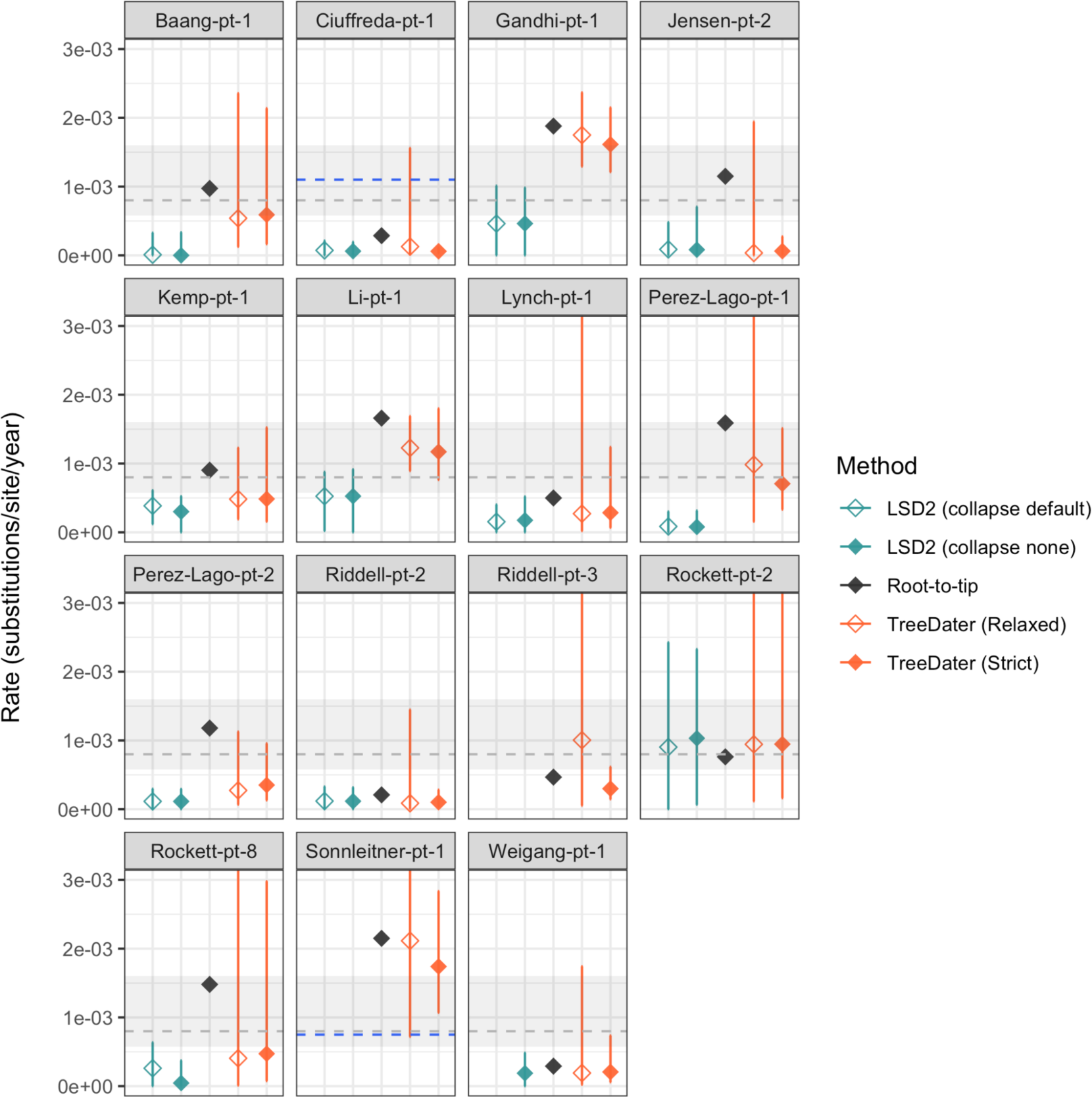
Substitution rate estimates for patients with ‘Questionable’ temporal signal. For all, rates were determined with following methods: LSD2 (with and without collapsing branches with short lengths), root-to-tip and TreeDater (by assuming relaxed and strict clock models). In each panel, the Y axis denotes the evolutionary rate in substitutions/site/year. For RTT only point estimates are represented. For other distance-based methods (i.e. LSD2 and TreeDater) diamonds represent mean estimates and horizontal lines illustrate confidence intervals. Rockett-pt-4 was removed as only RTT was successful (with rate estimate of 9.9e-03 substitutions/site/year). For Riddell-pt-3 and Sonnleitner-pt-1 evolutionary rate estimates could not be determined with LSD2. Similarly, for Weigang-pt-1 LSD2 analysis by assuming default node collapse value did not produce any results. Grey dashed line represents the commonly used SARS-CoV-2 substitution rate estimate of 8.00e-04 substitutions/site/year. The grey shaded area denotes the lowest and highest mean evolutionary rate estimates for SARS-CoV- 2 collected from various publications (5.75e-04 – 1.60e-03 subst./site/year, see Supplementary table S11). For Ciuffreda-pt-1 and Sonnleitner-pt-1 previously reported estimates of 1.1e-03 and 7.5e-4 substitutions/site/year, respectively, are indicated with a blue dashed line.

In figures 4 and 5 we compared the rates obtained within this study with three types of previously published estimates. Firstly, the grey dashed line represents a commonly used point estimate of 8.00e-04 substitutions/site/year reconstructed based on host-to-host transmission chains [57]. Secondly, the grey shaded area denotes the lowest and highest mean estimates collected from various publications describing evolutionary rates for SARS-CoV-2 host-to- host acute infections (5.75e-04 – 1.60e-03 subst./site/year, see Supplementary table S11). Thirdly, for those sample series for which a within-host rate was estimated in the source publication, this original estimate is indicated with a blue dashed line. This comparison revealed that for six out of nine patients with sufficient temporal signal the RTT estimate was higher than the point estimate of 8.00e-04 substitutions/site/year, whereas only for Harari-pt-5 the RTT estimate of 1.97e-03 exceeded all mean substitution rate estimates obtained from the literature (Figure 4). While some of the mean estimates from LSD2 or TreeDater analysis were higher than 8.00e-04, none of them exceeded the collection of mean estimates. However, for four of the sample series (Brandolini-pt-1, Halfmann-pt-1, Harari-pt-5 and Lee-pt-4) the widths of the confidence intervals revealed a considerable uncertainty in LSD2 and TreeDater estimates. Among these nine sample series, a previous intrahost rate estimate was available for Chaguza-pt-1. This estimate of 1.2e-03 substitutions/site/year was obtained with a root-to-tip regression approach and was therefore equal to our RTT estimate.

Figure 5 shows that for the majority of the datasets representing lower degrees of temporal signal the evolutionary rates obtained in this study were generally in good accordance with published host-to-host estimates: for most of the sample series the confidence intervals overlap with the grey shaded area representing the mean estimates from literature. Among these sample series with ‘Questionable’ temporal signal, within-host rates have been previously determined for Ciuffreda-pt-1 and Sonnleitner-pt-1. The reported rate of 0.09 mutations/day for Ciuffreda- pt-1 [32] translates into 1.1e-03 mutations/site/year, which is notably higher than estimates obtained in this study. On the contrary, the reported rate of 7.5e-4 substitutions/site/year for Sonnleitner-pt-1 [38], is considerably lower than RTT and TreeDater estimates derived in this study. However, the results should be interpreted cautiously since the genetic diversity and temporal spread of samples may not be sufficient to inform the molecular clock adequately.

### Evolutionary rate estimates – BEAST2

For the nine sample series exhibiting stronger temporal signals, evolutionary rates were also determined with BEAST v.2.6.7. The temporal signal, an essential prerequisite for Bayesian rate estimates [9,58], was additionally assessed with a date-randomization test (DRT) for these nine sample series. DRT results are presented in supplementary figures S3 and S4 for strict and uncorrelated relaxed clock models, respectively. Two criteria have been proposed for sufficient temporal signal in DRT results: 1) there is no overlap between posterior distributions of true and randomised [59], and 2) the true mean value is not contained in any of the randomised posterior distributions [19]. By assuming the Ramsden et al. 2009 criterion, DRT analysis of a strict clock model displayed strong evidence for sufficient temporal signal in four of the data series (Caccuri-pt-1, Chaguza-pt1, Halfmann-pt-1 and Khatamzas-pt-1). When assuming the more lenient criterion by Firth et al. 2010, datasets Choi-pt-1 and Lee-pt-4 were also included. Considering the uncorrelated relaxed lognormal clock model, strong temporal signal was observed only for Chaguza-pt-1 (Ramsden et al. 2009 criterion) or Chaguza-pt-1 and Khatamzas-pt-1 (Firth et al. 2010 criterion). For the rest of the sample series, as the 95% highest posterior density intervals (95% HPDIs) for the randomised datasets were somewhat overlapping with the real rate estimates, the strength of the temporal signal might not be sufficient to infer evolutionary rates with high confidence within a Bayesian framework.

Despite the DRT analysis not indicating a strong temporal signal particularly when assuming a relaxed clock model, evolutionary rates generated with BEAST2 were compared with estimates retrieved by other methods. For three sample series BEAST2 median estimates were in accordance with mean values obtained with LSD2 and TreeDater (Brandolini-pt-1, Huygens-pt-2 and Lee-pt-4), while for the rest of the sample series the median estimates inferred with BEAST2 were higher (Figure 4). Furthermore, for three sample series BEAST2 estimates exceeded the generally high RTT point estimates (Caccuri-pt-1, Choi-pt-1 and Khatamzas-pt-1). Overall, BEAST2 estimates showed less consistency than LSD2 and TreeDater relative to RTT. However, despite BEAST2 producing sporadically higher rates than other methods, similarly to RTT only Harari-pt-5 displayed a Bayesian median estimate exceeding the literature reference values used.

Whereas BEAST2 estimates obtained with strict and relaxed clock models were in good accordance within each sample series, evaluation of the estimated coefficient of rate variation alluded that for none of the nine sample series the evolutionary rate can be considered strictly constant through time (Supplementary figure S5). Although no precise criteria have been established in the literature, concentration of marginal posterior distribution of coefficient of rate variation below the value 0.1 can be considered sufficient to warrant the use of a strict clock model [60]. Thus, posterior distributions presented in Supplementary figure S5 support less clock-like evolution across branches of all nine datasets.

### Evaluating phylogenetic tree topologies and degrees of phylo-temporal clustering

To further examine the possible causes of the evolutionary rate estimate inconsistencies observed principally between BEAST2 vs. LSD2 and TreeDater, we inspected the topologies of SARS-CoV-2 phylogenetic trees. For each of the nine sample series topological distances between pairs of phylogenetic trees were calculated based on three comparisons: LSD2 vs. BEAST2 strict clock maximum clade credibility (MCC) tree, LSD2 vs. BEAST2 relaxed clock MCC tree, and BEAST2 strict clock MCC tree vs. BEAST2 relaxed clock MCC tree. Results are presented in Supplementary table S12. For the majority of the sample series, modest split differences were observed between LSD2 and both BEAST2 MCC trees. For two of the sample series, Caccuri-pt-1 and Huygens-pt-2, the score for conflicting splits exceeded the score for shared splits for comparisons between LSD2 vs. BEAST2 strict clock and LSD2 vs. BEAST2 relaxed clock. In contrast, tree topologies obtained with BEAST2 strict and BEAST2 relaxed clock models were identical for five of the sample series (Choi-pt-1, Halfmann-pt-1, Huygens- pt-2, Khatamzas-pt-1 and Lee-pt-4) and for the remaining four sample series only modest differences were detected between BEAST2 trees.

Further visual evaluation of the MCC trees generated with BEAST2 revealed a ‘ladder-like’ topology for the majority of the nine sample series (Supplementary figures S6–S14). This type of tree topology is considered indicative of excessive phylo-temporal clustering [61], which we further assessed by calculating temporal clustering (TC) statistics for all nine datasets. For four of the sample series – Chaguza-pt-1, Halfmann-pt-1, Harari-pt-5 and Khatamzas-pt-1 – we observed TC scores ranging between ∼0.3 and ∼0.5 (Supplementary table S13). Similar values have been interpreted to indicate a high degree of temporal clustering [62]. For these four datasets evolutionary rate estimates obtained with BEAST2 were notably higher than corresponding estimates produced with LSD2 or TreeDater. For Caccuri-pt-1 and Choi-pt-1 TC scores were less than 0.1, presumably indicating a lesser degree of phylo-temporal clustering. Whereas for Caccuri-pt-1 similar rate estimates were obtained with all methods, for Choi-pt-1 BEAST2 estimates are greater than LSD2 or TreeDater estimates. However, TC statistics are reported to be sensitive to small sizes below 20 [62] and thus a small sample size of nine sequences for Choi-pt-1 might affect its TC score. For the remaining three sample series (Brandolini-pt-1, Huygens-pt-2 and Lee-pt-4) we were not able to resolve the TC score unambiguously (for details, see Methods).

For Huygens-pt-2 a closer evaluation of MCC tree topologies (Supplementary figure 12), revealed significant substructure of the viral population. Whereas the first sequence for the sample series was obtained on the same day as the reported onset of symptoms (2022-01-06), the median estimates for the tree height date two months earlier with both clock models (2021- 11-07). Similar estimates for the most recent common ancestor were obtained with LSD2 (collapse none: 2021-11-03, collapse default: 2021-11-15), and TreeDater yielded even older estimates (strict clock: 2021-08-18, relaxed clock: 2021-09-17). Based on this, it is plausible that the patient has been superinfected with two SARS-CoV-2 strains representing the same Pango lineage (BA.1.1), and thus results for Huygens-pt-2 have been interpreted with caution.

### Patient case histories

We obtained evidence of non-clocklike evolution in nine sample series (Supplementary figure S5). Considering these, we were further interested in contrasting the timing of evolutionary rate changes with the temporal fluctuations in the viral load and the timing of SARS-CoV-2 treatments administered. As a proxy for viral load we used Ct values, information available for six of the patients (Brandolini-pt-1, Chaguza-pt-1, Choi-pt-1, Halfmann-pt-1, Harari-pt-5 and Huygens-pt-2). Additionally, direct estimates of viral load were given for Huygens-pt-2 and Khatamzas-pt-1. For these seven patients we additionally collected the SARS-CoV-2 treatment information, if any, from the original publications. Ct values, viral load estimates and timing points of SARS-CoV-2 treatments are given in Supplementary tables S2 and S4.

Changes in evolutionary rates through time were characterised by visualising MCC trees reconstructed with BEAST2 by assuming an uncorrelated relaxed clock model. However, as the BEAST2 estimates appeared biased towards higher rates, we further evaluated if the observed temporal oscillations in evolutionary rates hold when fixing the mean rate of relaxed clock model to a commonly used substitution rate reference estimate of 8.00e-04 substitutions/site/year. As shown in Supplementary figures S15–S21, when the inferred mean rate estimate is close to the fixed rate used, patterns of rate changes through time are highly similar between trees with fixed and unfixed clock rates (Brandolini-pt-1, Chaguza-pt-1 and Halfmann-pt-1). Conversely, when the inferred rate estimate is somewhat lower or higher than the fixed rate, minor scale differences can be detected between the corresponding trees (Choi- pt-1, Harari-pt-5, Huygens-pt-2 and Khatamzas-pt-1). Nonetheless, the broad patterns of evolutionary rate changes remain comparable, allowing for further examinations of temporal concurrencies.

‘Patient case histories’ for Chaguza-pt-1 and Khatamzas-pt-1, the only sample series for which temporal signal was adequate for relaxed clock analysis, are characterised in figures 6 and 7, respectively. For the rest of the sample series results are presented in Supplementary figures S22–S26. These seven patients displayed numerous different clinical conditions leading to a severely immunosuppressed condition (Table 1). Altogether the sample series covered a lengthy period of time from April 2020 to July 2022, during which new therapeutics for SARS-CoV-2 infection were developed. As a consequence, a notable variation in the treatment types can be detected among patients, antibody-based treatments targeting the spike protein – i.e. convalescent plasma, bamlanivimab, intravenous immunoglobulin and sotrovimab – being the most commonly used therapeutic agents. Two of the patients also received remdesivir with a direct antiviral activity targeting RNA polymerase. Moreover, the half-lives of different treatments vary greatly, ranging from a few hours for remdesivir [63,64] to nearly seven weeks for sotrovimab [65] (https://www.ema.europa.eu/en/medicines/human/EPAR/xevudy, last visited 20.10.2023). Convoluted cycling patterns of viral load were found in all nine patients, complicating a systematic comparison even further (Supplementary figure S27). While a visual examination revealed no strong evidence of temporal correspondences between molecular rate variation, viral loads, and the various SARS-CoV-2 treatments administered, adequate statistical testing was not possible due to the limited sample size and the reasons stated above.

**Figure 6.**
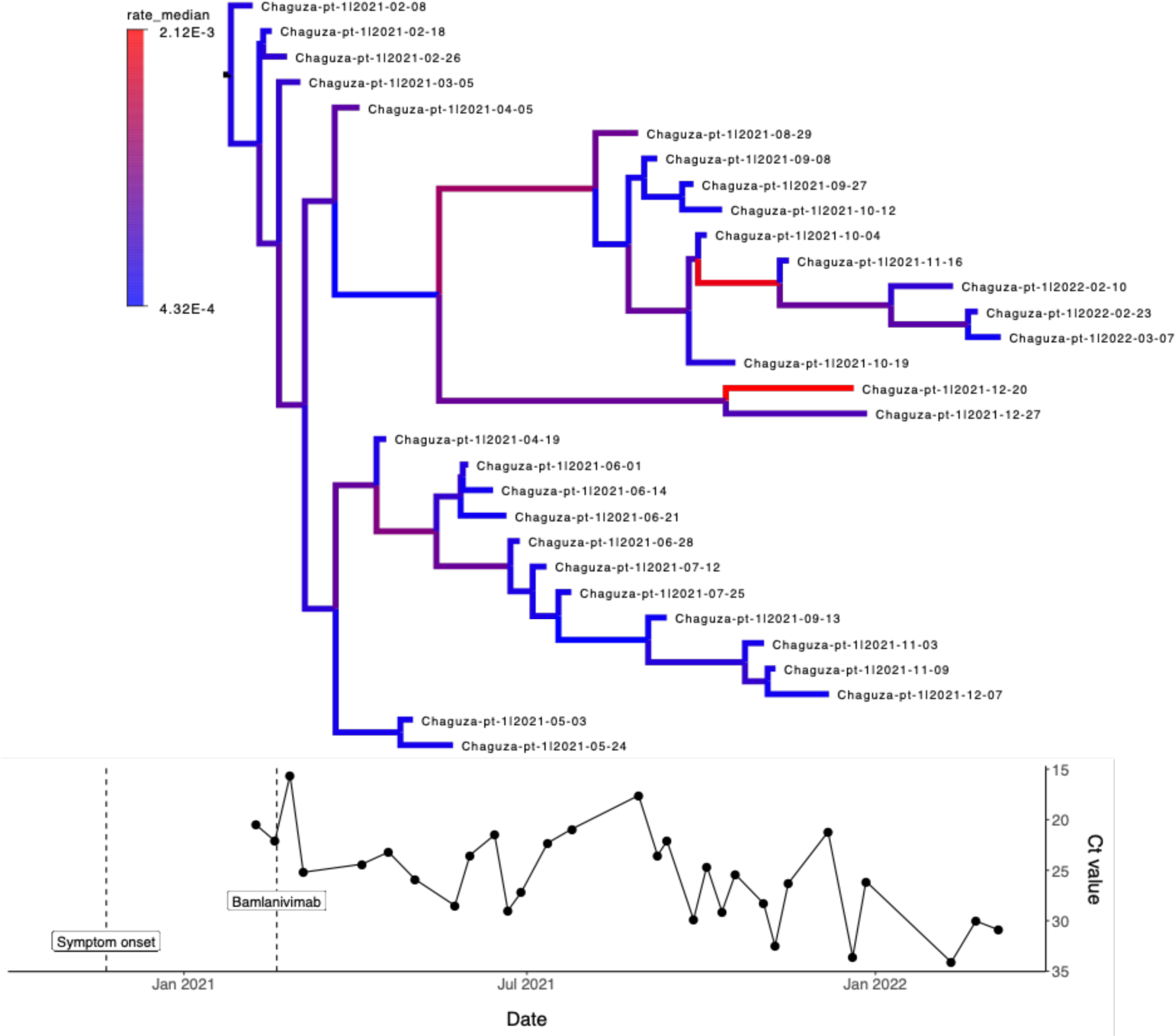
Patient case history for Chaguza-pt-1 patient, with advanced lymphocytic leukemia and B-cell lymphoma as underlying clinical conditions. Figure describes through time the changes in the evolutionary rates (by assuming an uncorrelated lognormal relaxed clock model), Ct values and SARS-CoV-2 treatments administered within the sampling window. For Chaguza-pt-1 sample series the first viral sequence was obtained 79 days after the onset of symptoms. Patient was treated with Bamlanivimab which targets spike-protein and has a half-time of approximately 17 days. Colouring of the branches within the phylogenetic tree represents evolutionary rate estimates (in substitutions/site/year) obtained with BEAST2, lower values indicated with blue and higher rates with red colour. Open circles denote samples for which only Ct values were available and coloured circles denote samples which were sequenced.

**Figure 7.**
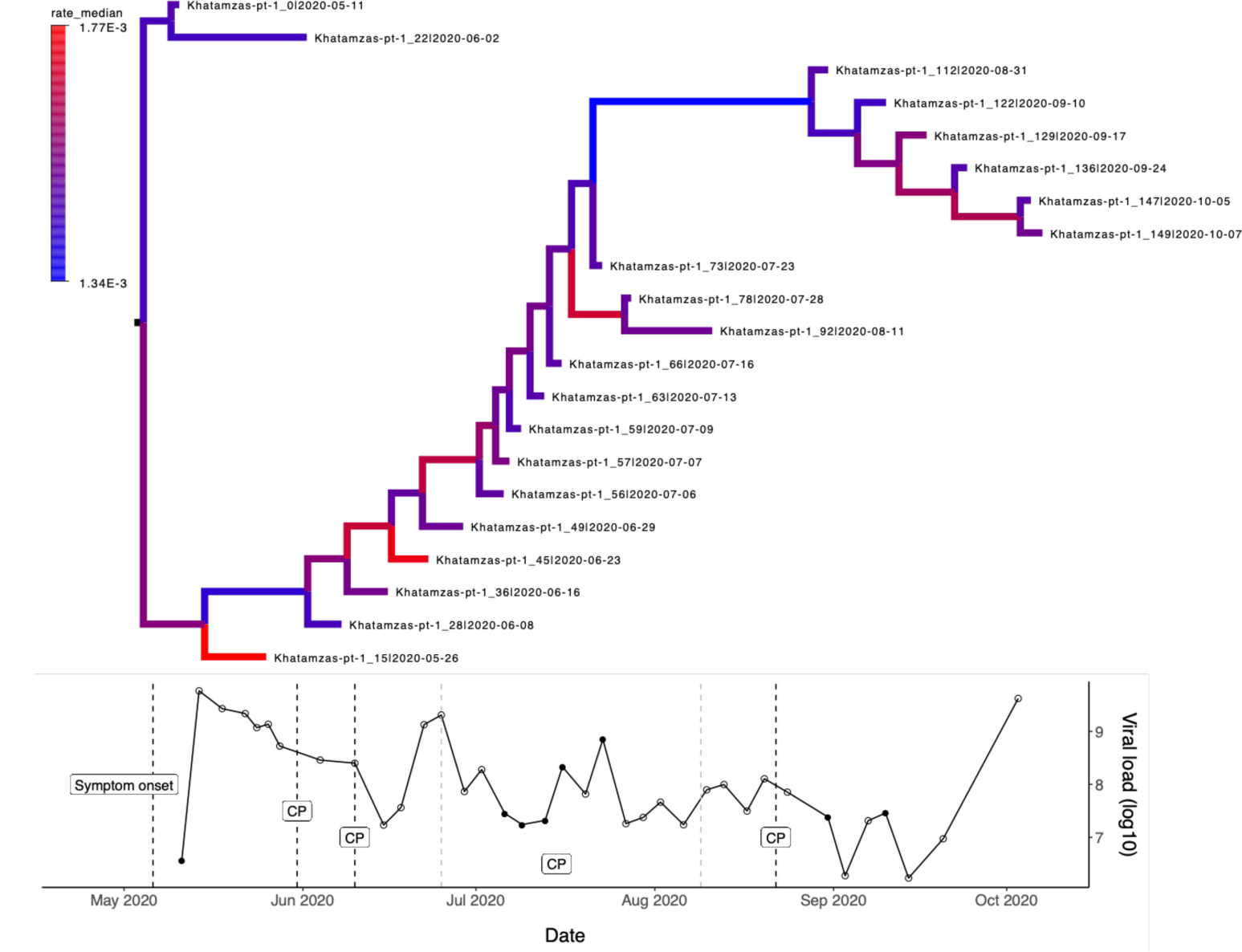
Patient case history for Khatamzas-pt-1 patient, with follicular lymphoma as underlying clinical condition. Figure describes through time the changes in the evolutionary rates (by assuming an uncorrelated lognormal relaxed clock model), viral load on a logarithmic scale and SARS-CoV-2 treatments administered within the sampling window. For the Khatamzas-pt-1 sample series the first viral sequence was obtained five days after the onset of symptoms. Patient was treated with convalescent plasma (CP) multiple times within the sampling window: on days 20, 30, 45-90 and 103 after the first sequenced sample (i.e. Day0). Convalescent plasma targets spike-protein and has a half-time of approximately 26 days with notable variation. Colouring of the branches within the phylogenetic tree represents evolutionary rate estimates (in substitutions/site/year) obtained with BEAST2, lower values indicated with blue and higher rates with red colour. For the viral load SARS-CoV-2 RNA copy numbers per ml of endotracheal aspirates are presented (See Khatamzas et al. 2022 Figure 1b) [50]. Open circles denote samples for which only viral load estimates were available and coloured circles denote samples which were sequenced.

## DISCUSSION

Chronic SARS-CoV-2 infections among immunocompromised individuals have been considered facilitative of an accelerated accumulation of mutations within a relatively short time window due to clinical conditions which are limiting the host’s immune response to the virus. However, most studies suggesting this lack the evaluation of temporal signal and the use of multiple methods of inference – two main principles for reliable tip-calibrated phylogenetic analyses. In this study we sought to fill in this gap by exploring intrahost dynamics of SARS-CoV-2 based on 26 viral sample series obtained from chronically infected individuals with a compromised immune system. The primary objective of this study is to evaluate the intrahost viral evolution from the molecular dating standpoint by inferring molecular rate estimates across the whole viral genome. We utilise a collection of commonly used phylogenetic approaches while simultaneously assessing the applicability and robustness of the methods and data utilised. In particular, our results exemplify the complexity of intrahost viral evolution.

### Low genetic diversity leading to insufficient temporal signal

The evaluation of within sample series’ genetic diversity revealed highly variable SARS-CoV- 2 diversity patterns among patients. Sample series showing lower genetic diversity despite long sampling window, such as Riddle-pt-3 with 225 days between first and last sample, conceivably indicate extremely low levels of viral replication for a lengthy period of time, as reported also in [66]. However, all 26 sample series included in this study exhibited genetic changes on a consensus sequence level over the course of infection. This is in contrast with the findings in [66], showing within-patient genetic variation in only around 30% of chronic infections. Differences between this study and [66] could be attributed to data discrepancies: whereas our dataset comprises viral sequences exclusively from patients with immunocompromised conditions, [66] included data from a large community-based surveillance study, likely containing individuals with a variety of clinical backgrounds including immunocompetent individuals. Nonetheless, given that no clinical metadata from [66] is publicly available, the true reasons for the observed disparities are unknown.

The low levels of genetic diversity observed were subsequently reflected in molecular dating analyses: whereas root-to-tip regression analysis suggested adequate temporal structure for all but one sample series (Lee-pt-11), a more rigorous evaluation through LSD2 and TreeDater analyses suggested sufficient temporal signal only for nine sample series. This exemplifies that RTT should be used only as an informal method for temporal signal assessment, as previously discussed for example in [9,10]. In addition, our results further confirm the previous statements proclaiming the problematic usage of root-to-tip regression as an explicit approach for molecular dating. Firstly, RTT assumes a strict clock model, whereas for all nine sample series for which rate heterogeneity was evaluated (through posterior distribution of the relaxed clock model’s rate parameter), the rate of evolution cannot be considered strictly constant through time. Secondly, even more severe biases might arise due to RTT’s simplified assumption of statistical independence of the sequences. The samples within the tree cannot be considered phylogenetically independent, instead they exhibit variable levels of shared ancestry. This leads to a pseudoreplication, where particularly the mutations occurring at the deeper branches of the tree are contributing to multiple root-to-tip distances. Supposedly sequences acquired from a prolonged intrahost infection are evolutionarily more closely related than a small collection of sequences randomly drawn from a large background population. This in turn will lead to more pronounced phylogenetic dependency for the intrahost sample series. As the RTT regression method accounts only for the absolute number of differences without explicitly modelling the shared ancestry of the sequences, estimates of the intrahost evolutionary rates can be highly inflated, as seen for the majority of the sample series included in this study. These varying degrees of phylogenetic dependence between a within-host and a population sample sets could potentially explain the notably higher intrahost evolutionary rates reported by Chaguza et al. 2023 and Stanevich et al. 2023 [30,36], as estimates were retrieved solely through root-to-tip regression analysis in these two studies.

### Notable variation in the rate estimates caused by the method-specific limitations

In order to evaluate whether previously reported accelerated intrahost evolutionary rates of SARS-CoV-2 can be seen as an artefact raised by the method applied, we exploited two additional distance-based methods: LSD2 and TreeDater. For the majority of sample series, the low phylogenetic signal produced high uncertainty in the parameter estimates resulting in wide confidence intervals seen particularly for TreeDater. In general, estimates generated using RTT tend to be consistently higher than rates obtained with LSD2 and TreeDater, which yielded rather similar results within each sample series. This applies also to the Chaguza-pt-1 sample series, for which RTT yielded a point estimate of 1.2e-03 substitutions/site/year (both this study and [30]) whereas LSD2 and TreeDater mean estimates were significantly lower, ranking from 4.6e-04 to 9.0e-04 substitutions/site/year. Mean estimates obtained with LSD2 and TreeDater were not overlapping with RTT 95% confidence interval reported in [30] (1.1e-03 – 1.3e-03 substitutions/site/year). This further supports our hypothesis of RTT introducing a noteworthy upward bias when employed on a dataset of evolutionary closely related sequences. It should be noted that the study by Stanevich et al. (2023), which reported within-host evolutionary rate of 1.53e-03 substitutions/site/year by utilising RTT method, was not part of this study. Sample series contained in total six sequences, two samples from August 2020 and four from January-February 2021, hence failing to meet our inclusion minimum of eight sequences. Despite data from Stanevich et al. 2023 was not included in this study, we would like to emphasise that rate estimations based upon small sample sizes with highly uneven temporal distribution should be interpreted with caution.

As a shared property of RTT, LSD2 and TreeDater is that they rely upon a user-specified substitution tree for which the optimal root position is estimated based on software specific algorithms. In the absence of a predefined outgroup, root estimation among genetically highly similar sequences can be challenging and may result in topological errors and biased rate estimates. To exclude the possibility of topological errors being the cause of the lower molecular rates obtained, we re-assessed rate estimates with LSD2 by utilising a SARS-CoV- 2 reference sequence as an outgroup. No significant differences were detected between the estimates reconstructed with and without an outgroup, suggesting that possible topological errors have only a modest impact on the inferred LSD2 rate estimates, if any, as also indicated by [11].

Despite molecular rate estimates being relatively robust for topological errors, we further exploited BEAST2 which, in contrast to distance-based methods, estimates probability distributions over parameters of interest, including the phylogenetic tree topology and evolutionary rate estimates. As Bayesian analyses have been considered rather sensitive to inadequate temporal signal [9,58] we chose to utilise only the nine sample series for which analysis with LSD2 and TreeDater suggested more discernible levels of temporal structure. Additional assessment of temporal signal through date-randomization test revealed that only for two of the sample series with the largest number sequences, Chaguza-pt-1 and Khatamzas- pt-1, accumulation of genetic diversity through time can be considered sufficient to allow the molecular rate to be inferred accurately with both strict and uncorrelated relaxed clock models (Table 2). For the rest of the datasets the strength of the temporal structure remains dubious particularly under the assumption of rate heterogeneity, suggesting that despite prolonged periods of infection somewhat low mutational rates of SARS-CoV-2 might not leave genetic signals strong enough for reliable molecular dating based on tip-calibration only.

**Table 2.**
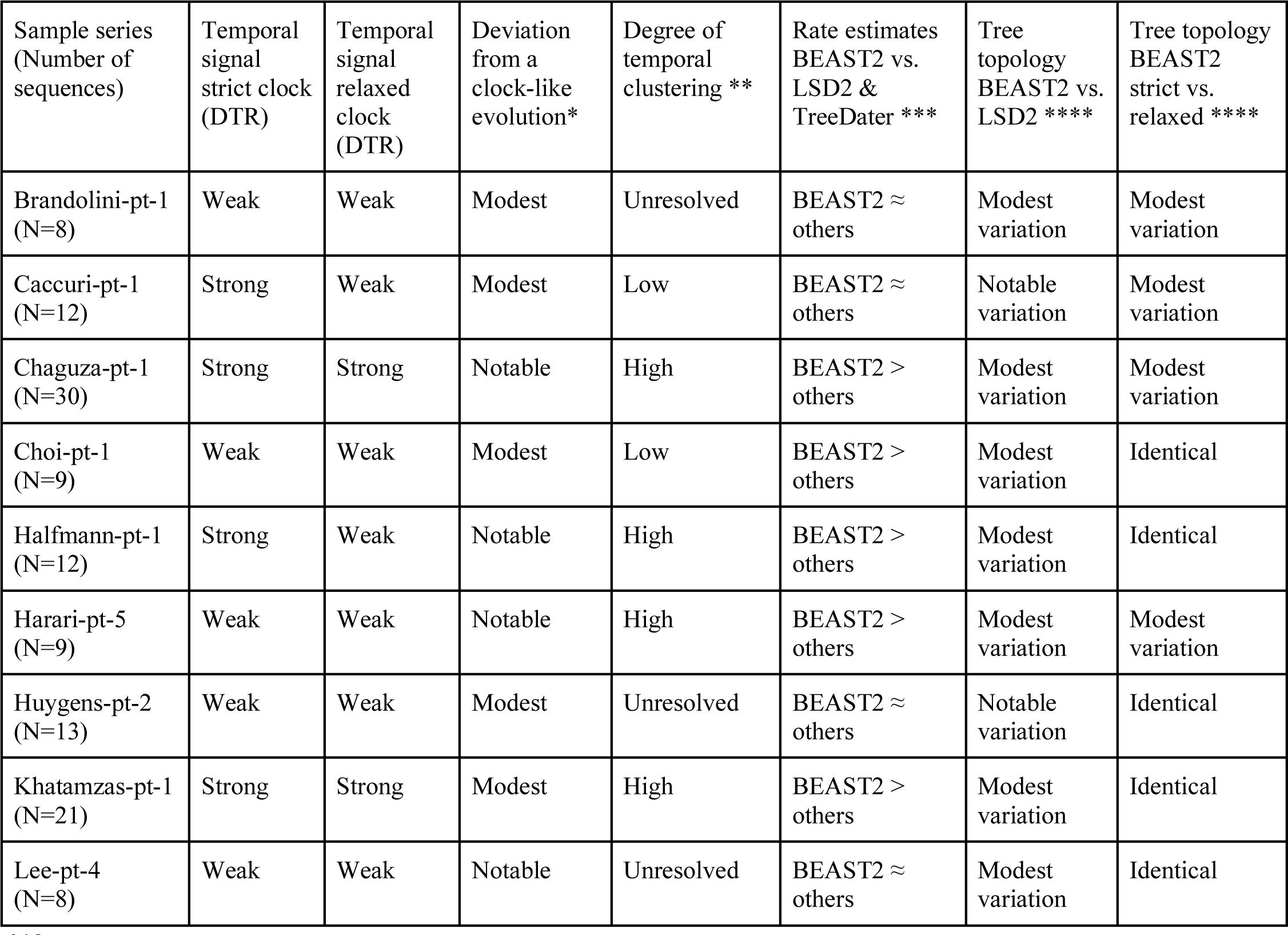
Summary of the results for nine sample series for which evolutionary rates were determined with LSD2, TreeDater and BEAST2. * Estimated based coefficient of rate variation (Supplementary figure S5). ** Estimated based on TC statistics (detailed values for three parallel runs are presented in Supplementary table S13). *** Comparison of point estimates (BEAST2 median estimates vs. LSD2 & TreeDater mean estimates) (see Figure 4 and Supplementary table S10). **** Estimated based on tree similarity and distance measures as proposed in Smith 2020 (detailed values presented in Supplementary table S12, see also Supplementary figures S6–S14).

In comparison to LSD2 and TreeDater results, rate estimates obtained using BEAST2 showed lesser degrees of consistency. We explored possible reasons for this variation by contrasting time-tree topologies generated with BEAST2 and LSD2 (Table 2, Supplementary table S12, Supplementary figures S6–S14). Whereas for some of the sample series distinctive structural disparities were observed between the trees, we couldn’t detect any systematic correlations between topological differences and inflated BEAST2 rate estimates explaining the variation (Table 2). A further evaluation of the underlying tree topology, however, revealed that the most plausible explanation for the high Bayesian rate estimates is temporal clustering of the samples. Ladder-like tree topology, where sequences obtained at similar times cluster together, tends to bear a strong phylo-temporal clustering [61]. Previous studies have demonstrated BEAST2 being profoundly sensitive to strong phylo-temporal clustering [58,67] as it decreases the number of independent calibration points, resulting in lower information content and increased uncertainty. This has been shown to yield an upward bias in Bayesian posterior estimates [67–69]. In contrast, LSD2 and TreeDater have shown to be less vulnerable for the presence of temporal clustering [11,13,67]. Although visual inspection of MCC trees indicated somewhat increased levels of phylo-temporal clustering for all nine sample series, reliable quantification of the temporal clustering statistic was only possible for the larger datasets, Chaguza-pt-1 and Khatamzas-pt-1. The clear indication of strong phylo-temporal clustering for Chaguza-pt-1 and Khatamzas-pt-1, plausibly explains high BEAST2 rate estimates for both sample series. Moreover, elevated levels of spatiotemporal clustering were presumably also reflected in poor convergence of the Bayesian analyses when uninformative clock rate prior was used (see Methods). Consequently, estimates derived with LSD2 and TreeDater are presumably closer to the true rates than those obtained with BEAST2.

For the Bayesian approach we chose to utilise as an underlying tree prior distribution a deterministic coalescent based Bayesian skyline plot (BSP) model [70] over the birth-death-sampling models. Despite the latter being considered more suitable for processes with stochastic population size changes including the emergence of a viral outbreak [71] modelling the within-host sampling process through time might be challenging, if not impossible. Given that poor characterization of the sampling process may lead to severely biased results within the birth-death-sampling framework [72] we considered a coalescent based approach being less vulnerable for misspecified sampling schemes. Moreover, we would like to point out that a comprehensive Bayesian analysis would also involve proper model selection to evaluate the best-fit clock and tree prior models, as well as sample-from-prior analysis, as discussed for example in [73,74]. However, given the vast number of sample series and various combinations of clock (strict vs. relaxed) and tree prior models (BSP vs. coalescent constant population size vs. coalescent exponential growth) to be tested, we chose to omit these further steps. Nonetheless, since misspecified tree prior may lead to increased rate estimates [75], we performed additional analyses for Chaguza-pt-1 and Khatamzas-pt-1 with coalescent constant size and exponential population growth models to ensure that elevated BEAST2 estimates are not a product of a tree prior used. Rate estimates inferred with these two additional tree prior models are greatly similar to estimates derived with BSP, as shown in Supplementary figure S28.

### Comparison with rate estimates obtained from acute infections provides no evidence for elevated intrahost rates

In previous studies, intrahost molecular rate estimates have been brought to a broader context through comparison with either 1) RTT estimates obtained from a randomly sampled background population [30,36] or with 2) a point estimate obtained from the literature [32,34,37]. In the latter case, the number of mutations accumulated is usually considered to directly reflect the within-host rate which is subsequently contrasted with a global rate estimate obtained at the early stages of the pandemic (i.e. ∼8.00e-04 substitutions/site/year). Later studies, however, have reported highly variable rates of SARS-CoV-2 evolution on a population scale, with mean estimates ranging from 5.75e-04 to 1.60e-03 substitutions/site/year [22], making inferences derived from a single point estimate somewhat ambiguous. Furthermore, the simplified assumption of genetic changes accumulating over a single viral lineage contradicts previous observations of chronic SARS-CoV-2 infection leading to the coexistence of genetically distinct viral populations, which could also be seen in some of the sample series included in this study (Brandolini-pt-1, Chaguza-pt-1, Huygens-pt-2, Khatamzas-pt-1).

Principally, a direct comparison of within-host and between-host rates may not be straightforward since molecular rate variation is not solely dependent on the rate of new mutations arising. Instead, the demographic history of the population has been found to alter the strength of genetic drift and selection, subsequently introducing rate variation through time [76–78] (for review see [79]). Patterns of rate variability have in addition shown to emerge due to ‘time-dependency’, proposing that molecular rate estimates rely on the length of the sampling window in question, with longer time intervals producing lower evolutionary rates [6,80]. Moreover, the degree of phylogenetic tree imbalance [81], the presence of a pronounced population structure [82] and the temporal distribution of sampling dates [83] have similarly been shown to impact the accuracy of inferred rate estimates. To mitigate the plausible biases introduced by demographic processes, we chose to compare rates derived in this study to a variety of previously published estimates which have been retrieved by using diverse methodologies and obtained from different datasets representing different timescales and phases of the pandemic (Supplementary table S11). Despite substantial discrepancies between sample series and methods used, intrahost evolutionary rates obtained in this study are generally consistent with rates reported from transmission chains of acutely infected individuals (Figure 4) and therefore our results do not support accelerated SARS-CoV-2 molecular rates within chronically infected immunocompromised individuals. Instead, our findings strongly suggest that within-host evolution across the whole SARS-CoV-2 genome is occurring at roughly the same rate as the background population.

### Fluctuations in the viral population size shaping the rate of molecular evolution?

A previous study by Chaguza et al. (2023) interpreted the elevated intrahost rates to reflect differences in viral population sizes. Unlike in host-to-host transmissions, the within-host pathogen population is not subject to transmission bottlenecks and thus intrahost SARS-CoV- 2 dynamics can result in faster evolutionary rates. However, since our results do not suggest notable differences between host-to-host and within-host rates, we further explored the possibility of changes in the viral population size leading to intrahost molecular rates comparable to estimates obtained from acute infections. Whereas serially sampled genealogies displaying excessive degrees of phylo-temporal clustering are traditionally thought to originate from viral populations under strong selective pressure [61], higher degrees of temporal clustering can also occur under neutral evolution as a result of repeated genetic bottlenecks [62]. Changes in the viral population size can be approximated, at least to some degree, either by directly expressing the amount of virus per unit volume of sample (i.e. viral load) or by test-specific cycle threshold (Ct) values, although both are sensitive to inconsistencies in sampling method (for a review see [26]). For the seven sample series with available data, frequent fluctuations in Ct or viral load estimates are apparent (Supplementary figure S27). Assuming that these reflect real changes in viral population sizes, these successive intrahost genetic bottlenecks might have caused a significant loss in genetic diversity, as shown also for example for *Staphylococcus aureus* [84]. Intriguingly, genetic diversity of intrahost respiratory tract samples – which comprised the majority of sample series used in this study – was found to be significantly lower when compared to other anatomic sites presumably leading to a more pronounced genetic drift [85]. Whereas the size of the intrahost genetic bottleneck is undoubtedly less stringent than what has been observed for host-to-host SARS-CoV-2 transmissions with one to 1000 viral particles transmitted between consecutive infections [24,86], repeated bottlenecks combined with a small effective population size and thus a greater impact of random sampling might temporarily affect the frequency of novel mutations emerging subsequently leading to lower molecular rates.

However, given that Ct values cannot be considered as a direct measurement of the viral population size, we further evaluated the signals of selection. Among nine of the sample series, only Lee-pt-4 showed evidence of positive selection across all functionally important proteins, albeit this was presumably driven by strong positive selection on ORF1ab which constitutes the vast majority of the SARS-CoV-2 genome (Supplementary table S14). For four of the sample series positive selection was detected on the S gene whereas the remaining four datasets showed no signals of selection. It is essential to note, however, that here the signals of the selection are tested by averaging over the entire length of a gene or genome. This implies that despite our findings not showing strong evidence of positive selection for entire genes or genomes, novel non-synonymous mutations, such as E484K and del144, have emerged and subsequently become fixed within sample series included in this study, indicating an excessive positive selection of individual antibody escape mutations. However, positive selection alone might be inefficient to produce the elevated levels of phylo-temporal clustering when accounting for the whole SARS-CoV-2 genome, leaving fluctuations in the population sizes as a plausible reason for the ladder-like tree topologies. As a result, we anticipate that intrahost population size variations can explain, at least to some extent, molecular rates analogous to host-to-host rates. Similar conclusions have been made for HIV in [87]. We acknowledge, however, the complexity of intrahost evolution of SARS-CoV-2. Whereas within-host population dynamics might partially explain the results observed in this study, more comprehensive understanding would require development of models accounting jointly for multiple evolutionary processes as discussed in [88] and as already available for instance for primary HIV infection [89].

### Complex patterns of non-clocklike evolution

As our findings indicate departures from the strictly clocklike evolution for all nine datasets investigated more thoroughly, we explored the possible factors causing episodic evolution through ‘Patient case histories’. Temporal correspondences of mutational patterns, viral loads and antibody-based treatments have previously been investigated, for example, by [41], where findings suggested strong evidence for a correlation between viral rebound and the emergence of antibody evasion mutations. We build upon the framework presented in [41], but instead of focusing on the emergence of individual mutations, our approach intends to explore mutational patterns on a more generic scale. Incorporating rate variation across branches could help us to comprehend evolutionary changes occurring between sampling points, providing insights into the general pace of viral evolution. This can provide information even for the unsampled parts of the phylogenetic tree. It is essential to note, however, that neither the approach used in this study nor the one exploited in [41] can reveal the exact timing of novel mutations emerging. More dense sampling over the course of infection would be required to be able to distinguish if certain antibody evasion mutations arose at the time of viral rebound or already during the preceding stages characterised by decreasing or undetectable levels of viral load. However, whereas proper statistical testing was not feasible due to numerous reasons (i.e. small sample size, complex cycling patterns of viral load and wide variation in clinical conditions as well as in SARS-CoV-2 treatments given) a visual examination of the ’Patient case histories’ does not explicitly reveal temporal simultaneity of viral rebound and elevated levels of viral evolution. Instead, our findings emphasise the complexities of the interplay between intrahost viral bottlenecks, molecular rate variation, and therapies targeting the virus, which are undoubtedly influenced by factors not explored here. Therefore, despite ‘Patient case histories’ being able to introduce an additional layer of information on intrahost viral evolution, larger cohorts and more samples as well as improved metadata documentation would be needed for statistically validated conclusions.

### Standardised framework for intrahost viral molecular rate inference needed

Whereas for SARS-CoV-2 the majority of molecular rate research has focused on rate variation at the population level, for other viruses, such as HIV, research on intrahost variation has been undertaken more extensively. Over the past three decades, a wide range of studies have reported within-host evolutionary rates for HIV (see for example references in Table 6 in [90]), intrahost estimates being consistently elevated compared to rates obtained from population scale phylogenies [91,92]. However, most of these estimates have been derived by depending solely on one dating method and, to the best of our knowledge, no systematic comparison of different approaches has been undertaken. Given that our findings clearly demonstrate the importance of comparing the results of multiple methods, we propose that studies estimating intrahost evolutionary rates of any virus could use the workflow established within this study. We recommend the following steps for robust tip-calibrated molecular dating inference of within-host sample series: 1) determination of genetic diversity, 2) evaluation of temporal signal, 3) exploration of the tree topology and 4) comparison of different molecular dating methods. This approach is particularly important when the phylogenies show strong signals of temporal clustering of the samples. We further note that the fundamental work established in [11,13,58,67,69], comparing distinct frameworks for molecular dating through simulation studies should be explored within intrahost datasets to gain more comprehensive understanding of method specific limitations.

### Limitations of the study – Sequence analysis related constraints

Our study has several limitations, the most prominent of which arise from the data itself. Despite the fact that we utilised a collection of 26 sample series, the date-randomisation test showed that the lack of phylogenetic signal hampered adequate assessment of molecular rates for all except two of the largest datasets (Chaguza-pt-1 and Khatamzas-pt-1). While our data inclusion criteria of at least eight sequences was arbitrarily chosen, our findings suggest that with lower numbers of sampled genomes all the genetic variants and thus the entire viral diversity may not be well represented and temporal differences of the evolutionary response may go undetected. However, minimum sample size used in this study should not be referred to as a generally recognised threshold, instead each dataset’s eligibility for molecular dating analysis should be assessed individually. Furthermore, we utilised consensus sequences as provided by the original publications implying that distinct methodologies as well as different variant calling thresholds have been used for consensus sequence reconstruction among sample series (see Supplementary table S6 for details). However, since we mainly focus on molecular dating method comparison on a within sample series level, possible biases introduced by differences between consensus sequence reconstruction methods can be considered negligible. As a more general complication it should be noted that when utilising only consensus sequences single nucleotide variants (SNPs) prevalent at low frequencies are ignored and the data do not represent the full genetic diversity of the intrahost viral population, as shown for example in [85]. However, based on our literature and database searches, the availability of raw sequence data in public repositories is even more restricted than what is seen for consensus sequences. As a further limitation can be seen that we chose to derive rate estimates for the entire SARS-CoV-2 genome, as is standard practice for both inter- and intra-host sample series. However, studies have reported evolutionary rate variation between different genomic regions [93,94], leaving the characterisation of gene-specific rate differentiations between within-host and host-to-host viral evolution an open question for future research. Despite occurrences of intrahost recombination of two distinct viral variants being reported [95,96] we didn’t explore the possibility of prolonged SARS-CoV-2 infection facilitating recombination either 1) between intrahost viral variants and lineages circulating in the background population or 2) between coexisting within-host quasispecies. To minimise the possibility of sequences being recombinants of two different viral variants we required as an inclusion criteria evidence in the original publication confirming the occurrence of a long-term infection (i.e. not multiple independent infections or superinfection) and further verified that all sequences within a sample series represented the same Pango lineage. For the latter, the possible recombination events are likely to remain undetected due to the high consensus sequence similarity of coexisting quasispecies and the low overall genetic diversity resulting in too few polymorphic sites for reliable recombination analysis.

### Limitations of the study – Metadata-related constraints

Despite a large number of published SARS-CoV-2 sequences collected from immunocompromised patients globally, our finding that only 26 individuals had a series of at least eight sequences available demonstrates the relative scarcity of high-resolution genetic analyses. Furthermore, we observed a substantial degree of variation in data collection practices, which serves to hinder the direct comparison of multiple datasets. Moreover, as we show here, the viral phylogenies are not sufficient alone to inform us on within-host dynamics of SARS-CoV-2. Instead, joint analysis of multiple non-mutually exclusive processes, including the host’s immune system, viral population dynamics and administered treatments, is required to understand the underlying drivers shaping the phylogenetic tree. Through our exploration of ‘Patient case histories’ we develop a framework for simultaneously evaluating both genetic and clinical datasets. We thus propose that samples, as well as associated patient metadata, should be collected systematically over the course of infection. To model better the interplay between genetic drift and adaptive selection, one would need metadata that characterises viral population size changes (i.e. viral load estimates or Ct values) as well as information on factors possibly impacting the selection (i.e. information on underlying clinical conditions, treatments and vaccination status). Whereas the overwhelming majority of sequences used in this study were derived from either oropharyngeal or nasopharyngeal swabs (258/323), the lack of gastrointestinal or serological specimens collected reveals an important underexploited avenue of research. Indeed, consistent practices of sampling multiple tissue types would likely be informative for our understanding of intrahost disease dynamics including viral reservoirs, which have been hypothesised to play a role within long COVID [97], impacting millions of people and causing a huge economical. We believe that the imposition of minimum standards for metadata collection, as well as the incentivisation and enforcement of data sharing will be important steps in facilitating improved interdisciplinarity in the future.

### Conclusions

Our findings have two types of implications: firstly, the results of this study emphasise the complexity of determining the within-host evolutionary rates, not restricted only to intrahost evolution of SARS-CoV-2 but generalised also for other pathogens. By neglecting the limitations of the data or the method used, it is possible to derive highly biased rate estimates and to draw invalid conclusions. Our findings highlight the significance of conducting a systematic study of several sample series using different approaches in order to support reliable estimations. In the absence of previously established standards, we propose that future studies estimating within-host viral molecular rates could follow, when applicable, the workflow established within this study. Secondly, in terms of SARS-CoV-2, our findings provide no evidence of greater levels of viral evolution in immunocompromised patients with chronic SARS-CoV-2 infection when considering the complete viral genome. Instead, within-host molecular rates are comparable with rate estimates derived from host-to-host transmission chains not restricted to immunocompromised individuals. While our findings challenge previous claims of increased intrahost evolutionary rates, they do not refute the generally recognised theory of immunocompromised individuals serving as a source for emergence of new viral variants. Whereas for the sample series included in this study the intrahost evolution likely proceeds at a rate similar to that of the background population, a prolonged SARS-CoV- 2 infection within an immunodeficient patient might promote the appearance of novel antibody escape mutations. Furthermore, our findings do not preclude the possibility of increased evolutionary rates among immunocompromised individuals, however, no viral data from such a chronic infection was identified within this study.

## MATERIALS AND METHODS

### Data collection

All data used within this study was obtained through a literature search conducted between 15.08.2022 - 15.03.2023, according to the search terms: *Case study; longitudinal; SARS-CoV- 2; COVID; immunocompromised; persistent; prolonged; viral evolution; intra-host; long-term*. The resulting dataset of 1,029 longitudinally sampled consensus sequences from 255 patients and 53 publications was then filtered according to the following criteria: (i) given evidence within the original publication of the immunocompromised status of the individual, (ii) confirmation that the infection was the result of a single, long-term infection, i.e. excluding multiple consecutive infections, or a superinfection, (iii) that at least 8 sequences with unique collection dates were available from the patient, with the aim of minimising phylogenetic uncertainty and thus increasing the precision of parameter estimates. We furthermore followed the procedure presented in Harari et al 2022 and considered an individual to have a chronic SARS-CoV-2 infection if there was evidence of persistent viral shedding for a period of at least 20 days. The removal of all patients not fulfilling these criteria resulted in a final dataset of 323 consensus sequences from 26 patients and 21 publications. For the sample series obtained from [52], the last sample (EPI_ISL_2484152, 2020-07-08) was excluded from all the analyses since in the original publication authors suspected a superinfection with a second strain of the virus.

In parallel to sequence data collection, clinical metadata obtained from the original publications or via correspondence with the authors are provided within supplementary tables S1-S6. For consistency, all sample series were renamed according to the first author of the source publication, followed by ‘pt’ and the patient number. This labelling is used throughout the manuscript and the original patient identifiers are listed in supplementary table S7. Sequence identifiers were renamed according to the day of collection, where in each case ‘day 0’ represented the earliest sequence available for the patient. In some instances, multiple samples were collected on the same day, representing different specimen types (e.g. Baang-pt-1_22a and Baang-pt-1_22b). In such cases, only one sample was considered for a given collection date and preference was given to respiratory tract samples, since within-host populations from different tissue types have been shown to be genetically highly distinctive [85]. Pango lineages were obtained from original publications and were further confirmed with Nextclade v2.14.1 [98].

### Genetic diversity

Sequences were aligned to the SARS-CoV-2 reference genome (NC_045512.2) in MAFFT v7.475 [99] with the --keeplength option. Within the group mean number of pairwise differences were determined with MEGA 11 [100]. Distances were estimated by calculating the absolute number of differences by assuming uniform rates among sites and treating gaps and missing data as pairwise deletion. As a variance estimation method we assumed bootstrap with 100 replications.

### Evolutionary rate estimates – RTT, LSD2 and TreeDater

For each sampling series, consensus sequences were aligned as described previously and alignment ends as well as other possibly problematic positions were masked, as suggested in [101]. For each sample series we assessed the strength of temporal signal with root-to-tip linear regression with the R package BactDating [102]. For BactDating, the input substitution trees were generated with IQ-Tree v2.1.2 [103] simultaneously estimating the best-fit substitution model with ModelFinder [104] (iqtree2 -s input.fasta -m MFP). At this point, the temporal signal was considered sufficient for the downstream analysis if the p-value of R^2^ was less than 0.05.

Subsequently, for sample series with RTT confirmed temporal signal, evolutionary rate estimates were assessed with Least-Squares Dating (LSD2) method integrated in IQ-TREE v2.1.2 as well as with. For both methods, the maximum likelihood substitution tree inferred with IQ-Tree was provided as an input. Time trees were inferred by using sampling dates as tip dates and the root position was estimated as a part of the analyses. For LSD2 the best-fit substitution model was estimated with ModelFinder, as described previously. Regarding the tree we chose to use two different approaches: Within the first approach we followed the LSD2 default values and collapsed all internal branches having branch length less than 1.67e-05 (= 0.5/sequence length). Within the second approach, none of the branches were collapsed implying that null branches were allowed. For the output tree branch lengths were resampled in total 100 times to determine the confidence intervals (with *--date-ci* option). With TreeDater the molecular rates were determined by assuming a strict and relaxed clock. For both, confidence intervals for the rate estimates were estimated with a parametric bootstrap with 100 replicates.

Based on the results obtained from LSD2 and TreeDater, the strength of temporal signal of each sample series was re-evaluated: If LSD2 and/or TreeDater analysis yielded error messages (see below) indicating a poor temporal signal, the temporal signal for the sample series under scrutiny was considered as ‘Questionable’. The software specific error messages considered were:

1. LSD2: The estimated rate reaches the given lower bound (1e-10).
2. TreeDater: Warning: Root to tip regression predicts a substitution rate less than zero.

Tree may be poorly rooted or there may be small temporal signal.

### Evolutionary rate estimates – BEAST2

For the sample series passing the re-evaluation of the temporal signal the evolutionary rates were additionally determined with BEAST v.2.6.7. Evolutionary rates were inferred with strict and uncorrelated relaxed lognormal clock models by assuming a Bayesian Skyline Plot (BSP) as an underlying tree model. Due to small sample sizes, dimensions for BSP model parameters bPopSize and bGroupSize were set to 3–5, depending on the data set. As a substitution model HKY + Γ was used. As a prior distribution for a strict clock rate parameter (clockRate) an uniform distribution (0,1) was used. Same uniform distribution of (0,1) was originally used also for relaxed clock rate parameter (ucldMean). However, Markov Chain Monte Carlo (MCMC) chains were not reaching convergence. Therefore, we chose to use more stringent prior and set normal distribution with mean of 0.0008 and standard deviation of 0.0016. No additional modifications were made to the default prior distributions. The temporal signal was assessed with a date-randomization test (DRT) implemented in R package TIPDATINGBEAST [105]. For the DRT, for each sample series for both clock models 20 randomised data sets were generated as recommended in [69].

The MCMC chain length was set to 10–50 million steps for all MCMC analyses. For real data analysis the posterior distributions of parameters were estimated based on two parallel MCMC chains. After confirming the sufficient convergence of each chain (effective sample sizes for each parameter > 200), the samples from two runs were combined after discarding the first 10% of each chain as a burn-in. Maximum clade credibility trees with median node heights were reconstructed with TreeAnnotator by assuming 10% as a burn-in. MCC trees were visualised with FigTree v1.4.4 (http://tree.bio.ed.ac.uk/software/figtree/, last visited 20.10.2023).

### Estimating topological distances

The topological distances between pairs of phylogenetic trees were estimated with the R package ‘TreeDist’ v.2.6.3 [106] (https://zenodo.org/records/3528124, last visited 26.10.2023), which is an information-based generalised Robinson-Foulds metric that defines the overall similarity between two trees. For each sample series three comparisons were performed: LSD2 vs. BEAST2 strict clock MCC tree, LSD2 vs. BEAST2 relaxed clock MCC tree, and BEAST2 strict clock MCC tree vs. BEAST2 relaxed clock MCC tree. According to [106], ‘SharedPhylogeneticInfo’ metrics describes the amount of phylogenetic information in common between two trees, whereas ‘DifferentPhylogeneticInfo’ metrics describes the distance between trees under scrutiny i.e. how much information is different in the splits of these two trees. Regarding LSD2, comparisons were performed with allowing zero length branches and collapsing short branches. Results for the latter are presented in parenthesis. When ‘DifferentPhylogeneticInfo’yielded a value of 0, trees were considered identical. When the score for shared splits exceeded the score for conflicting splits (‘SharedPhylogeneticInfo’ > ‘DifferentPhylogeneticInfo’), two trees were considered to exhibit modest variation in the tree topology. When the score for conflicting splits exceeded the score for shared splits (‘SharedPhylogeneticInfo’ < ‘DifferentPhylogeneticInfo’), trees were considered to exhibit notable variation in the tree topology.

### Evaluating the degree of phylo-temporal clustering

The degree of temporal clustering was estimated by calculating temporal clustering (TC) statistics [62] implemented in R package PhyloTempo [107]. As an input, we used the same unrooted substitution trees generated with IQ-Tree which we used as input also for BactDating, LSD2 and TreeDater. For each sample series the TC score was defined with three independent runs by setting the number of randomizations to 500. In case these three separate analyses produced highly divergent TC score estimates, we considered the degree of temporal clustering as unresolved.

### Test of positive selection

The presence of positive selection was evaluated through a codon-based Z-test of selection averaging over all sequence pairs within the dataset for nine of the sample series. As a null hypothesis we assumed a strict-neutrality (dN = dS) and as an alternative hypothesis positive selection (dN > dS). All calculations were conducted with MEGA 11 [100] by using Pamilo- Bianchi-Li method by assuming a pairwise deletion.

## AUTHOR CONTRIBUTIONS

Conceptualization: Sanni Översti, Emily Gaul, Denise Kühnert Data curation: Emily Gaul, Sanni Översti, Björn-Erik O. Jensen Formal analysis: Sanni Översti, Emily Gaul

Funding acquisition: Denise Kühnert Investigation: Sanni Översti, Emily Gaul

Methodology: Sanni Översti, Emily Gaul, Denise Kühnert Project administration: Sanni Översti, Emily Gaul Resources: Björn-Erik O. Jensen

Validation: Sanni Översti, Emily Gaul Visualization: Sanni Översti, Emily Gaul

Writing – original draft: Sanni Översti, Emily Gaul

Writing – review & editing: Sanni Översti, Emily Gaul, Denise Kühnert, Björn-Erik O. Jensen

## ACKNOWLEDGEMENTS

We wish to sincerely acknowledge the valuable contribution of all those authors who provided us further information regarding their data. We gratefully thank the following researchers: Dr. Carlos Flores, Dr. Laura Ciuffreda, MSc. Jose M. Lorenzo-Salazar, Dr. Julia Alcoba-Florez, MD Sammy Huygens, Dr. Bart Rijnders, Dr. Xiaoli Wang, Dr. Rebecca Rockett, Dr. Vitali Sintchenko, Dr. Sissy T. Sonnleitner, Dr. Eva Hinterbichler and Dr. Gernot Walder. Moreover, the authors wish to thank Gerd Specht for his technical assistance with PhyloTempo analysis. We also thank Dr. Jukka Palo for his insightful remarks on the manuscript draft. We gratefully acknowledge all data contributors, i.e., the Authors and their Originating laboratories responsible for obtaining the specimens, and their Submitting laboratories for generating the genetic sequence and metadata and sharing via the GISAID Initiative, on which this research is based.

## DATA AVAILABILITY

No new data was created as a part of this study. Instead, the findings in this study are based on previously published datasets. For the majority of the sample series included, accession information for the viral genomic sequences is given in the Supplementary table S3. Additionally, viral genomic data generated for Sonnleitner et al. 2022, is available in the Genome Sequence Archive as .bam files under the bioproject name PRJCA008906 (https://ngdc.cncb.ac.cn/bioproject/browse/PRJCA008906). Corresponding consensus sequences for can be obtained through correspondence with the authors of Sonnleitner et al. 2022. Viral genomic data generated for Li et al. 2021 can be obtained through correspondence with the authors of Li et al. 2021. Viral genomic data generated for Jensen et al. 2021 can be obtained through correspondence with the authors of Jensen et al. 2021. Files associated with phylogenetic analysis will be available in GitHub: https://github.com/tidelab/Chronic_covid_evolutionary_rates.

## SUPPORTING INFORMATION

**Supplementary table S1.** Sequence metadata.

**Supplementary table S2.** Patient metadata.

**Supplementary table S3.** Sequence accession information.

**Supplementary table S4.** All Reported Ct Values / Viral Loads of Patient Viral Specimens.

**Supplementary table S5.** List of supporting publications.

**Supplementary table S6.** Information on bioinformatics procedures used in each supporting publication.

**Supplementary table S7.** Patient list.

**Supplementary table S8.** Nextstrain clades, Pango lineages and WHO variant of concern (VOC) statuses for sample series included in this study.

**Supplementary table S9.** Mean number of pairwise differences between sequence pairs. See also Figure 3.

**Supplementary table S10.** Evolutionary rate estimates reconstructed with RTT, LSD2, TreeDater and BEAST2. Evolutionary rates are given in substitutions/site/year. For LSD2 and TreeDater mean estimates are given with lower and upper bounds of confidence intervals. For estimates inferred with BEAST2 median estimates with 95% highest posterior density intervals (HPDI) are presented.

**Supplementary table S11.** Evolutionary rates obtained from literature and used as a reference. Table is an extension to the table presented in Attwood et al. 2022 (Table 1). Abbreviations as in Attwood et al. 2022: BCP = Bayesian coalescent phylodynamic, MTBD = Multi-type birth-death, SC = Structured coalescent, BC + EG = Bayesian coalescent with exponential growth.

**Supplementary table S12. Topological distances between pairs of phylogenetic trees.** For each sample series, three comparisons were performed with R package ‘TreeDist’: LSD2 vs. BEAST2 strict clock MCC tree, LSD2 vs. BEAST2 relaxed clock MCC tree, and BEAST2 strict clock MCC tree vs. BEAST2 relaxed clock MCC tree. According to Smith 2020, ‘SharedPhylogeneticInfo’ metrics describes the amount of phylogenetic information in common between two trees, whereas ‘DifferentPhylogeneticInfo’ metrics describes the distance between trees under scrutiny i.e. how much information is different in the splits of these two trees. Regarding LSD2, comparisons were performed with allowing zero length branches and collapsing short branches. Results for the latter are presented in parenthesis. When ‘DifferentPhylogeneticInfo’yielded a value of 0, trees were considered identical. When the score for shared splits exceeded the score for conflicting splits (‘SharedPhylogeneticInfo’> ‘DifferentPhylogeneticInfo’), two trees were considered to exhibit modest variation in the tree topology. When the score for conflicting splits exceeded the score for shared splits (‘SharedPhylogeneticInfo’<‘DifferentPhylogeneticInfo’), trees were considered to exhibit notable variation in the tree topology (highlighted with red colour).

**Supplementary table S13.** Results from the PhyloTempo analysis performed for nine sample series. The temporal clustering (TC) statistics can get values between 0 and 1, TC=0 indicating a complete absence of temporal clustering (Gray et al. 2011, Nordström et al. 2012). In Gray et al. 2011 TC values of ∼ 0.3 and above are considered to indicate a high degree of TC. ‘Staircase-ness’ statistic describes the proportion of imbalanced subtrees (Nordström et al. 2012) and values of zero indicate a perfectly balanced binary tree whereas values of one indicate a perfectly imbalanced tree (Nordström et al. 2012). Degree of temporal clustering was considered as ‘Unresolved’ for those sample series for which TC scores obtained from three independent runs were highly divergent. Under the TC scores, the optimal number of time intervals as well as number of leaves assigned to each bin, are reported for each parallel run.

**Supplementary table S14.** Results from Z-test of positive selection. Table cells represent the test statistic (dN - dS) and green colour demonstrates statistically significant indication of positive selection (i.e. p values < 0.05). * All = ORF1ab, S, E, M and N.

**Supplementary figure S1.** Root-to-tip regression plots for 25 sample series included in this study (Lee-pt-11 omitted due to lack of temporal signal). R package BactDating was used to perform regression of root-to-tip analysis and to generate the figures. Note, that as BactDating requires sampling dates in calendar units, for sample series lacking collection dates in calendar years (i.e. Baang-pt-1, Gandhi-pt-1, Jensen-pt-2 and Kemp-pt-1) the collection day for Day0 sequence was arbitrarily set to 2020-01-01 and collection dates for the rest of the sequences were calculated accordingly (for example for Baang-pt-1: Day5 sample → 2020-01-06, Day15 sample → 2020-01-16, etc.). Therefore, for these four sample series the timescales on the x axis do not indicate the actual sampling window.

**Supplementary figure S2.** Testing the impact of inclusion of an outgroup for evolutionary rate estimates inferred with LSD2. As an outgroup reference sequence NC_045512.2 was used. In each panel, the Y axis denotes the evolutionary rate in substitutions/site/year. Diamonds represent mean estimates obtained without an outgroup (i.e. the best-fit root position is estimated according to LSD criteria) whereas triangles represent mean estimates obtained when a tree is being rooted with a known outgroup. Grey dashed line represents the commonly used SARS-CoV-2 substitution rate estimate of 8.00e-04 substitutions/site/year. The grey shaded area denotes the lowest and highest mean evolutionary rate estimates for SARS-CoV-2 collected from various publications (5.75e-04 – 1.60e-03 subst./site/year, see Supplementary table S11).

**Supplementary figure S3.** Date-randomisation test (DRT) performed on clockRate parameter of the strict clock model. Each panel corresponds to estimates obtained from one sample series. Within each panel, an estimate indicated with red colour represents the real estimate whereas black colour denotes estimates obtained from date-randomized data sets. For each sample series date-randomization was performed twenty times. For clarity, on the Y axis evolutionary rate estimates are reported on a logarithmic scale. Overlapping 95% highest posterior density (HPD) distributions of real and randomized estimates might indicate that the strength of the temporal signal might not be sufficient enough to infer evolutionary rates with high confidence only based on tip-dating.

**Supplementary figure S4.** Date-randomisation test (DRT) performed on ucldMean parameter of the uncorrelated relaxed lognormal clock model. Each panel corresponds to estimates obtained from one sample series. Within each panel, an estimate indicated with red colour represents the real estimate whereas black colour denotes estimates obtained from date- randomized data sets. For each sample series date-randomization was performed twenty times. For clarity, on the Y axis evolutionary rate estimates are reported on a logarithmic scale. Overlapping 95% highest posterior density (HPD) distributions of real and randomized estimates might indicate that the strength of the temporal signal might not be sufficient enough to infer evolutionary rates with high confidence only based on tip-dating.

**Supplementary figure S5.** Marginal posterior distributions for coefficient of rate variation (uncorrelated lognormal relaxed clock model). This parameter characterises the clock-likeness of the data, and values closer to zero suggest that a strict clock model might describe the data better. Whereas no rigorous value threshold has been given in literature, most often the usage of strict clock model is considered justified when majority of the probability mass is placed below 0.1 (indicated with red dashed line). Posterior distributions for all nine sample series illustrate signals of non-clocklike evolution, rate variation among branches being pronounced especially in Chaguza-pt-1, Halfmann-pt-1, Harari-pt-5, Khatamzas-pt-1 and Lee-pt-4.

**Supplementary figure S6.** Time-trees for Brandolini-pt-1. In the upper panel maximum clade credibility (MCC) trees from BEAST2 strict (left) and relaxed (right) clock analysis are given. In the lower panel a maximum likelihood tree generated with LSD2 is given. For the LSD2 tree, internal branches having branch length less than 1.67e-05 (= 0.5/sequence length) were collapsed. For BEAST2 trees node posterior support values are presented, for LSD2 bootstrap values.

**Supplementary figure S7.** Time-trees for Caccuri-pt-1. In the upper panel maximum clade credibility (MCC) trees from BEAST2 strict (left) and relaxed (right) clock analysis are given. In the lower panel a maximum likelihood tree generated with LSD2 is given. For the LSD2 tree, internal branches having branch length less than 1.67e-05 (= 0.5/sequence length) were collapsed. For BEAST2 trees node posterior support values are presented, for LSD2 bootstrap values.

**Supplementary figure S8.** Time-trees for Chaguza-pt-1. In the upper panel maximum clade credibility (MCC) trees from BEAST2 strict (left) and relaxed (right) clock analysis are given. In the lower panel a maximum likelihood tree generated with LSD2 is given. For the LSD2 tree, internal branches having branch length less than 1.67e-05 (= 0.5/sequence length) were collapsed. For BEAST2 trees node posterior support values are presented, for LSD2 bootstrap values.

**Supplementary figure S9.** Time-trees for Choi-pt-1. In the upper panel maximum clade credibility (MCC) trees from BEAST2 strict (left) and relaxed (right) clock analysis are given. In the lower panel a maximum likelihood tree generated with LSD2 is given. For the LSD2 tree, internal branches having branch length less than 1.67e-05 (= 0.5/sequence length) were collapsed. For BEAST2 trees node posterior support values are presented, for LSD2 bootstrap values.

**Supplementary figure S10.** Time-trees for Halfmann-pt-1. In the upper panel maximum clade credibility (MCC) trees from BEAST2 strict (left) and relaxed (right) clock analysis are given. In the lower panel a maximum likelihood tree generated with LSD2 is given. For the LSD2 tree, internal branches having branch length less than 1.67e-05 (= 0.5/sequence length) were collapsed. For BEAST2 trees node posterior support values are presented, for LSD2 bootstrap values.

**Supplementary figure S11.** Time-trees for Harari-pt-5. In the upper panel maximum clade credibility (MCC) trees from BEAST2 strict (left) and relaxed (right) clock analysis are given. In the lower panel a maximum likelihood tree generated with LSD2 is given. For the LSD2 tree, internal branches having branch length less than 1.67e-05 (= 0.5/sequence length) were collapsed. For BEAST2 trees node posterior support values are presented, for LSD2 bootstrap values.

**Supplementary figure S12.** Time-trees for Huygens-pt-2. In the upper panel maximum clade credibility (MCC) trees from BEAST2 strict (left) and relaxed (right) clock analysis are given. In the lower panel a maximum likelihood tree generated with LSD2 is given. For the LSD2 tree, internal branches having branch length less than 1.67e-05 (= 0.5/sequence length) were collapsed. For BEAST2 trees node posterior support values are presented, for LSD2 bootstrap values.

**Supplementary figure S13.** Time-trees for Khatamzas-pt-1. In the upper panel maximum clade credibility (MCC) trees from BEAST2 strict (left) and relaxed (right) clock analysis are given. In the lower panel a maximum likelihood tree generated with LSD2 is given. For the LSD2 tree, internal branches having branch length less than 1.67e-05 (= 0.5/sequence length) were collapsed. For BEAST2 trees node posterior support values are presented, for LSD2 bootstrap values.

**Supplementary figure S14.** Time-trees for Lee-pt-4. In the upper panel maximum clade credibility (MCC) trees from BEAST2 strict (left) and relaxed (right) clock analysis are given. In the lower panel a maximum likelihood tree generated with LSD2 is given. For the LSD2 tree, internal branches having branch length less than 1.67e-05 (= 0.5/sequence length) were collapsed. For BEAST2 trees node posterior support values are presented, for LSD2 bootstrap values.

**Supplementary figure S15**. Impact of fixing the mean rate of relaxed clock analysis for Brandolini-pt-1. In the left, mean rate is estimated with prior N(0.0008, 0.0016) and in the right mean rate is fixed to 8.00e-04 substitutions/site/year.

**Supplementary figure S16.** Impact of fixing the mean rate of relaxed clock analysis for Chaguza-pt-1. In the left, mean rate is estimated with prior N(0.0008, 0.0016) and in the right mean rate is fixed to 8.00e-04 substitutions/site/year.

**Supplementary figure S17.** Impact of fixing the mean rate of relaxed clock analysis for Choi- pt-1. In the left, mean rate is estimated with prior N(0.0008, 0.0016) and in the right mean rate is fixed to 8.00e-04 substitutions/site/year.

**Supplementary figure S18.** Impact of fixing the mean rate of relaxed clock analysis for Halfmann-pt-1. In the left, mean rate is estimated with prior N(0.0008, 0.0016) and in the right mean rate is fixed to 8.00e-04 substitutions/site/year.

**Supplementary figure S19.** Impact of fixing the mean rate of relaxed clock analysis for Harari-pt-5. In the left, mean rate is estimated with prior N(0.0008, 0.0016) and in the right mean rate is fixed to 8.00e-04 substitutions/site/year.

**Supplementary figure S20.** Impact of fixing the mean rate of relaxed clock analysis for Huygens-pt-2. In the left, mean rate is estimated with prior N(0.0008, 0.0016) and in the right mean rate is fixed to 8.00e-04 substitutions/site/year.

**Supplementary figure S21.** Impact of fixing the mean rate of relaxed clock analysis for Khatamzas-pt-1. In the left, mean rate is estimated with prior N(0.0008, 0.0016) and in the right mean rate is fixed to 8.00e-04 substitutions/site/year.

**Supplementary figure S22.** Patient case history for Brandolini-pt-1 patient, with follicular lymphoma as underlying clinical condition. Figure describes through time the changes in the evolutionary rates (by assuming an uncorrelated lognormal relaxed clock model), Ct values and SARS-CoV-2 treatments administered within the sampling window. For Brandolini-pt-1 the first viral sequence was obtained 132 days after the onset of symptoms. Patient was treated with intravenous immunoglobulin (IVIG) which targets spike-protein and has a half-time of approximately 26 days with notable variation. Colouring of the branches within the phylogenetic tree represents evolutionary rate estimates (in substitutions/site/year) obtained with BEAST2, lower values indicated with blue and higher rates with red colour. Open circles denote samples for which only Ct values were available and coloured circles denote samples which were sequenced.

**Supplementary figure S23.** Patient case history for Choi-pt-1 patient, with catastrophic antiphospholipid syndrome (CAPS) as underlying clinical condition. Figure describes through time the changes in the evolutionary rates (by assuming an uncorrelated lognormal relaxed clock model), Ct values and SARS-CoV-2 treatments administered within the sampling window. For Choi-pt-1 the first viral sequence was obtained 18 days after the onset of symptoms. Patient was treated twice with Remdesivir which targets polymerase and has a half-time of approximately 17 hours. Patient was also treated with an antibody cocktail against SARS-CoV-2 (Regeneron, Baum et al. 2020). Colouring of the branches within the phylogenetic tree represents evolutionary rate estimates (in substitutions/site/year) obtained with BEAST2, lower values indicated with blue and higher rates with red colour. Open circles denote samples for which only Ct values were available and coloured circles denote samples which were sequenced.

**Supplementary figure S24.** Patient case history for Halfmann-pt-1 patient, with primary immunodeficiency as underlying clinical condition. Figure describes through time the changes in the evolutionary rates (by assuming an uncorrelated lognormal relaxed clock model), Ct values and SARS-CoV-2 treatments administered within the sampling window. For Halfmann- pt-1 the first viral sequence was obtained 113 days after the onset of symptoms. Patient was treated with multiple SARS-CoV-2 treatments within the sampling window. Colouring of the branches within the phylogenetic tree represents evolutionary rate estimates (in substitutions/site/year) obtained with BEAST2, lower values indicated with blue and higher rates with red colour. Open circles denote samples for which only Ct values were available and coloured circles denote samples which were sequenced.

**Supplementary figure S25.** Patient case history for Harari-pt-5 patient, with acute lymphoblastic leukemia (ALL) as underlying clinical condition. Figure describes through time the changes in the evolutionary rates (by assuming an uncorrelated lognormal relaxed clock model), Ct values and SARS-CoV-2 treatments administered within the sampling window. For Harari-pt-5 the first viral sequence was obtained on the same day as the onset of symptoms. Patient was treated with convalescent plasma (CP) in total four times: on days 33 & 34 and 42 & 43 after the onset of symptoms. Convalescent plasma targets spike-protein and has a half-time of approximately 26 days with notable variation. Colouring of the branches within the phylogenetic tree represents evolutionary rate estimates (in substitutions/site/year) obtained with BEAST2, lower values indicated with blue and higher rates with red colour. Open circles denote samples for which only Ct values were available and coloured circles denote samples which were sequenced.

**Supplementary figure S26.** Patient case history for Huygens-pt-2 patient, with lymphoma as underlying clinical condition. Figure describes through time the changes in the evolutionary rates (by assuming an uncorrelated lognormal relaxed clock model), Ct values and SARS-CoV- 2 treatments administered within the sampling window. For Huygens-pt-2 the first viral sequence was obtained on the same day as the onset of symptoms. Patient was treated with Sotrovimab, which targets the spike-protein and has a half-time of approximately 49 days. Additionally, the patient was treated with convalescent plasma (CP). Colouring of the branches within the phylogenetic tree represents evolutionary rate estimates (in substitutions/site/year) obtained with BEAST2, lower values indicated with blue and higher rates with red colour. Open circles denote samples for which only Ct values were available and coloured circles denote samples which were sequenced.

**Supplementary figure S27.** Ct values (upper panel) and viral load (lower panel) for seven of the sample series. Open circles denote samples for which only Ct values were available and coloured circles denote samples which were sequenced. For Huygens-pt-2 both Ct values and viral load estimates were available.

**Supplementary figure S28.** Rate estimates for Chaguza-pt-1 and Khatamzas-pt-1 obtained with alternative tree priors. For the results presented in the main text, for BEAST2 analysis the Bayesian skyline plot (BSP) model was used as an underlying tree prior. For sample series Chaguza-pt-1 and Khatamzas-pt-1 we performed additional analysis by assuming coalescent constant size and coalescent exponential population growth models. For both tree priors, runs were executed by assuming strict and uncorrelated lognormal relaxed clock models. Results show that tree priors do not have a notable impact on the evolutionary rate estimates inferred.

